# Influence of DNA sequences on thermodynamic and structural stability of ZTA transcription factor - DNA complex: An all-atom molecular dynamics study

**DOI:** 10.1101/2024.11.03.621784

**Authors:** Boobalan Duraisamy, Debabrata Pramanik

**Affiliations:** Department of Physics, SRM University AP, Amaravati 522 240, Andhra Pradesh, India; Centre for Computational and Integrative Sciences, SRM University AP, Amaravati 522 240, Andhra Pradesh, India

**Keywords:** Epstein–Barr virus, ZTA Transcription factor, protein-dsDNA complex, ZTA responsive elements, molecular dynamics, umbrella sampling, free energy, fingerprint calculation

## Abstract

The Epstein-Barr virus (EBV) is one of the cancer-causing gamma type viruses. Although more than 99% people are infected by this virus at some point, it remains in the body in a latent state, typically causing only minor symptoms. Our current understanding is that a known transcription factor (TF), the ZTA protein, binds with dsDNA (double stranded deoxyribonucleic acid) and plays crucial role in mediating the viral latent-to-lytic cycle through binding of specific ZTA responsive elements (ZREs). However, there is no clear understanding of the effect of DNA sequences on the structural stability and quantitative estimation of the binding affinity between the ZTA TF and DNA, along with their mechanistic details. In this study, we employ integrated classical all-atom molecular dynamics (MD) and enhanced sampling simulations to study the ZTA-dsDNA structural properties, thermodynamics, and mechanistic details for the ZTA protein and for two different dsDNA systems: core motif and core motif with flanking end sequences. For each system, we studied three different ZTA responsive elements (ZREs) sequences: ZRE 1, ZRE 2 and ZRE 3. We performed structural analyses, including RMSD and RMSF calculations, to assess conformational stability, along with detailed interaction profiles and hydrogen bond analysis. We conducted residue-level and nucleic acid-level analyses to assess the important protein residues and DNA bases forming interactions between the ZTA and dsDNA systems. We also explored the effect of adding flanking end sequences to the core motif on DNA groove lengths and interstrand hydrogen bonds. Our results indicate that the flanking sequences surrounding the core motif significantly influence the structural stability and binding affinity of the ZTA-dsDNA complex. Among ZRE 1, ZRE 2, and ZRE 3, particularly when paired with their naturally occurring flanking ends, ZRE 3 exhibits higher stability and binding affinity. These findings provide insights into the molecular mechanisms underlying EBV pathogenesis and may indicate potential targets for therapeutic intervention. A detailed of the binding mechanisms will allow for the design of better-targeted therapies against EBV-associated cancers. This study will serve as a holistic benchmark for future studies on these viral protein interactions.

## INTRODUCTION

The gamma-type Epstein-Barr virus (EBV) is the first identified human tumour virus, responsible for promoting various diseases including multiple sclerosis, nasopharyngeal carcinoma^1^, breast cancer^2^, gastric cancer^3^, hepatitis^4^, and more. It affects populations globally, with an infection rate of more than 90%^5^, and the mortality rate has been reported to be comparatively higher in East Asian countries^6^. It transmits through body fluids,^7,8^ such as saliva, sweat, and physical contact. Infected individuals exhibit symptoms such as fever, enlarged tonsils, and sore throat^7^. Diagnostic tests like PCR (polymerase chain reaction) and in-situ hybridization can identify antibodies generated during the acute phase of infection^9,10^. The life cycle of this virus is biphasic with latent and lytic phases. Within memory B cells, the dormant virus resides during the latent phase without replication, causing persistent infection^11^. The histopathological findings reveal the presence of EBV infection in biopsy specimens of infected patients^11–14^. Under certain conditions, such as immune suppression, the dormant virus can transition to the lytic phase, leading to active replication and production of virions^15,16^.

The latent to lytic switch is orchestrated by ZTA or Zebra, from the family of basic leucine zipper transcription factors (bZIP-TF). ZTA is a pivotal protein encoded during the initial phase of infection by the EBV IE (Epstein-Baar virus immediate early) gene BZLF1 (BamHI Z fragment leftward open reading frame 1). It can activate early and late genes by binding with viral dsDNA^17–20^. ZTA comprises 245 amino acids (aa) and is structured into five distinct domains: the transcriptional activation domain (aa 1-167), the putative regulatory domain (aa 168-177), the DNA binding domain (aa 178-194), the coil-coil domain (aa 195-227), and the accessory activation domain (aa 228-245)^21,22^. Among these five domains, we studied the ZTA TF residues ranging from 178 – 236, which are reported in the crystal structure (PDB ID: 2C9N)^23^. The complete genome sequence of Epstein Barr virus is available under NCBI reference ID: NC_007605.1^24^ and has been analysed to facilitate various studies^25,26^.

The ZTA TF–DNA complex plays a major role in latent-to-lytic activation. Therefore, understanding^27–29^ the binding nature of ZTA TF with DNA, the effect of mutations in ZTA, the substitution of nucleic acids in the dsDNA core-motif region, and the role of DNA flanking ends, is of paramount importance to explore. Several previous studies have revealed that ZTA prefers binding to methylated DNA (me-C) binding sites over ZREs. This selective affinity is attributed to the presence of the Serine 186 residue in the DNA-binding domain^20,30,31^. Mutations within the DNA-binding domain of ZTA result in functional alterations and modulate its interaction with viral dsDNA. The S186A mutation in ZTA reduces DNA binding affinity, particularly to methylated sites, impeding the transition of the lytic cycle and downstream gene expression induction^20,31–33^. Earlier, Lee Heston *et al.* carried out targeted mutations across all the residues in the DNA binding domain (178 to 194) and reported mutations that enhance binding affinity alongside those exhibiting impaired binding capabilities^34^. Later, Ray *et al.* elucidated the effects of mutations at the Cystein-189^35^ and Asparagine-182^36^ positions with various substitutions, revealing diverse impacts on DNA binding specificity. Further studies revealed that mutations in the DNA-binding domain at R179A, Y180E, and A185V not only influence the lytic switch but also disrupt the activation of EA-D (the viral DNA polymerase processivity factor)^37^. Prior works investigated the interaction between the ZTA protein and viral dsDNA, focusing on unique sequences of core motifs. We have studied three different sequences: ZRE 1 (TGAGCCA), ZRE 2 (TGAGCGA), and ZRE 3 (TTCGCGA),^32^ respectively. These ZRE sequences have been taken from the Viral GENOME sequence shown in Figure 1. Previous experimental studies reported that changes in the core motif sequences in dsDNA affects the DNA binding affinity. Yella et al. observed that alterations in the flanking sequences significantly impact the binding affinity of the protein–DNA systems^38^. Although several experimental studies have reported the ZTA–DNA interaction and its functional intricacies, there are still no studies providing a quantitative estimation of the binding affinity between the ZTA TF and various core–motif DNA sequences, the effect of DNA flanking ends added to the core motif, and mechanistic insights behind the ZTA-dsDNA interaction.

**Figure 1:**
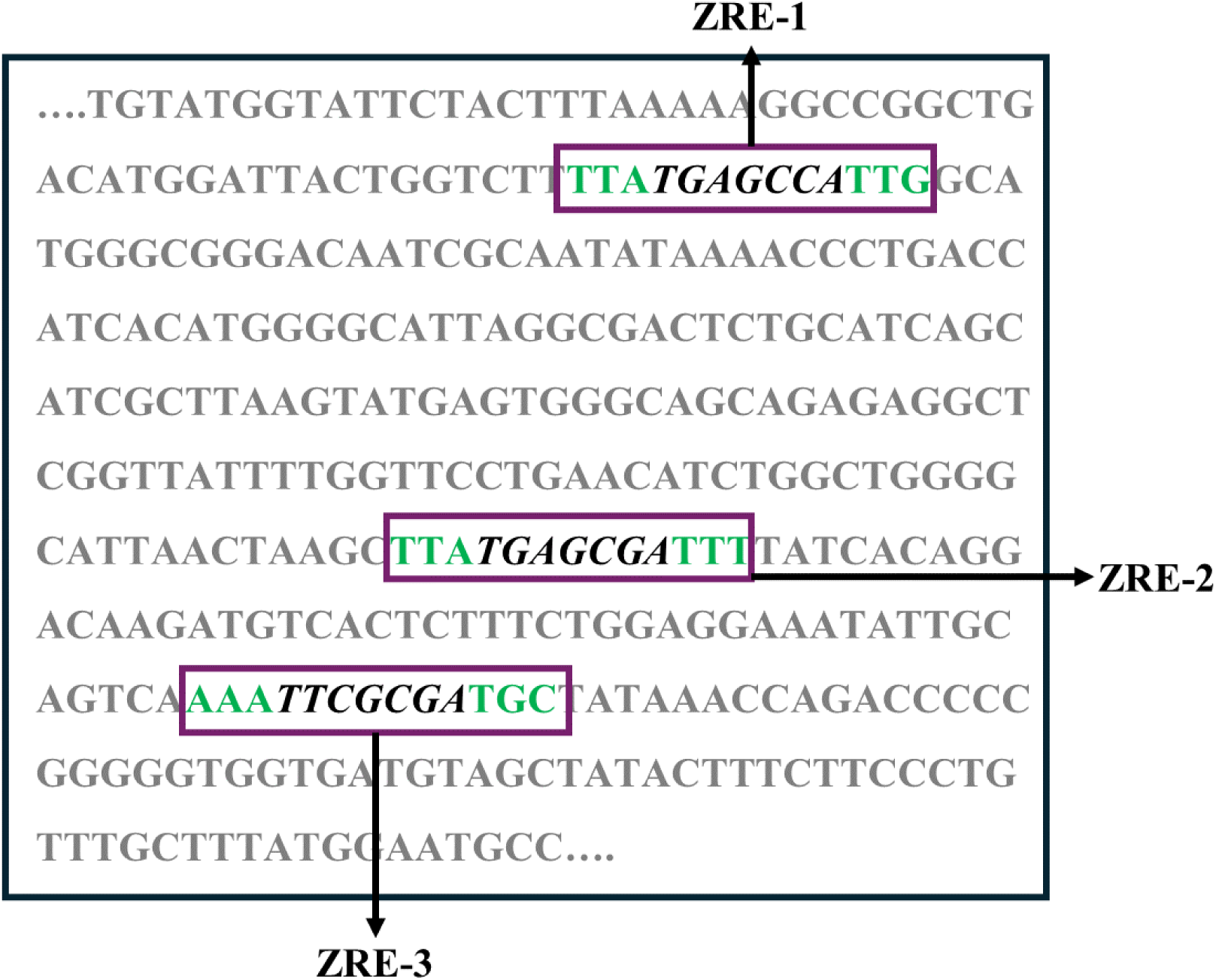
ZRE 1, ZRE 2 and ZRE 3 sequences are shown from available viral genome sequences reported in NCBI reference ID: NC_007605.1^24^. In this figure we have shown core motif sequences in the highlighted box in black colour for different ZRE’s whereas the corresponding flanking end sequences are marked in green colour.

Previous studies have shown that molecular dynamics (MD) simulations can be a major tool in exploring the structural properties^14,39–41^ thermodynamic behaviour^42–45^, interactions, and microscopic picture of complex systems like protein – ligand interactions^46–52^, protein – DNA interactions^27–29^, and DNA – nanomaterials ^43,53–56^

To understand the complete picture of the ZTA interaction with viral dsDNA at the molecular level, we employed classical all-atom MD simulations integrated with the umbrella sampling method^57–59^ and studied the thermodynamics of the ZTA-DNA interaction. In 2006, Carlo Petosa et al. published the crystal structure of ZTA bound with viral dsDNA (PDB ID-2C9N, 2C9L)^23^, and more recently, Bernaudat et al. presented the ZTA crystal structure with PDB ID- 7NX5^30^. In this work, we conducted computational simulations for the ZTA-DNA system, focusing on all ZREs for the core motifs and core motifs with flanking end sequences as reported in NCBI reference ID: NC_007605.1^24^. We performed a comparative study across all categories to assess the binding affinities and structural dynamics of the ZTA-DNA interactions in varying contexts and observed variations in the results for each category. Understanding the dynamics between ZTA and viral DNA is paramount for deciphering the molecular mechanisms underlying the Epstein-Barr virus pathogenesis. It not only explains the regulatory roles of ZTA in viral gene expression but also offers valuable insights into potential therapeutic targets for combating EBV-associated diseases.

## COMPUTATIONAL METHODOLOGY

### Structural modelling

The ZTA-dsDNA structure was modelled using the crystal structure available in the PDB (Protein data bank) database (PDB ID:2C9N). The missing atoms and residues in the ZTA protein–dsDNA crystal structure were modelled using Swiss-Modeler^60,61^. In this work, six different sequences of dsDNA have been studied (provided in Table 1). These new modified sequences in DNA were developed using the UCSF Chimera tool^62^. Discovery studio^63^ was utilized to superimpose structures onto the original crystal structure for analysis purposes (Figure 2(a)). We performed molecular dynamics (MD) simulations at the all-atom level using Gromacs-2022.4^64^ as the basic MD engine. We parametrized the ZTA protein using Amber03 force field parameters and dsDNA using Amber 94 parameters, respectively^65^. To model the aqueous environment, the TIP3P^66^ water model was used. The system was charge neutralized by adding Na+ and Cl- counterions to substitute for the water molecules in the system. To mimic the bulk environment, we applied periodic boundary conditions (PBC) in all three directions of the simulation box.

**Figure 2:**
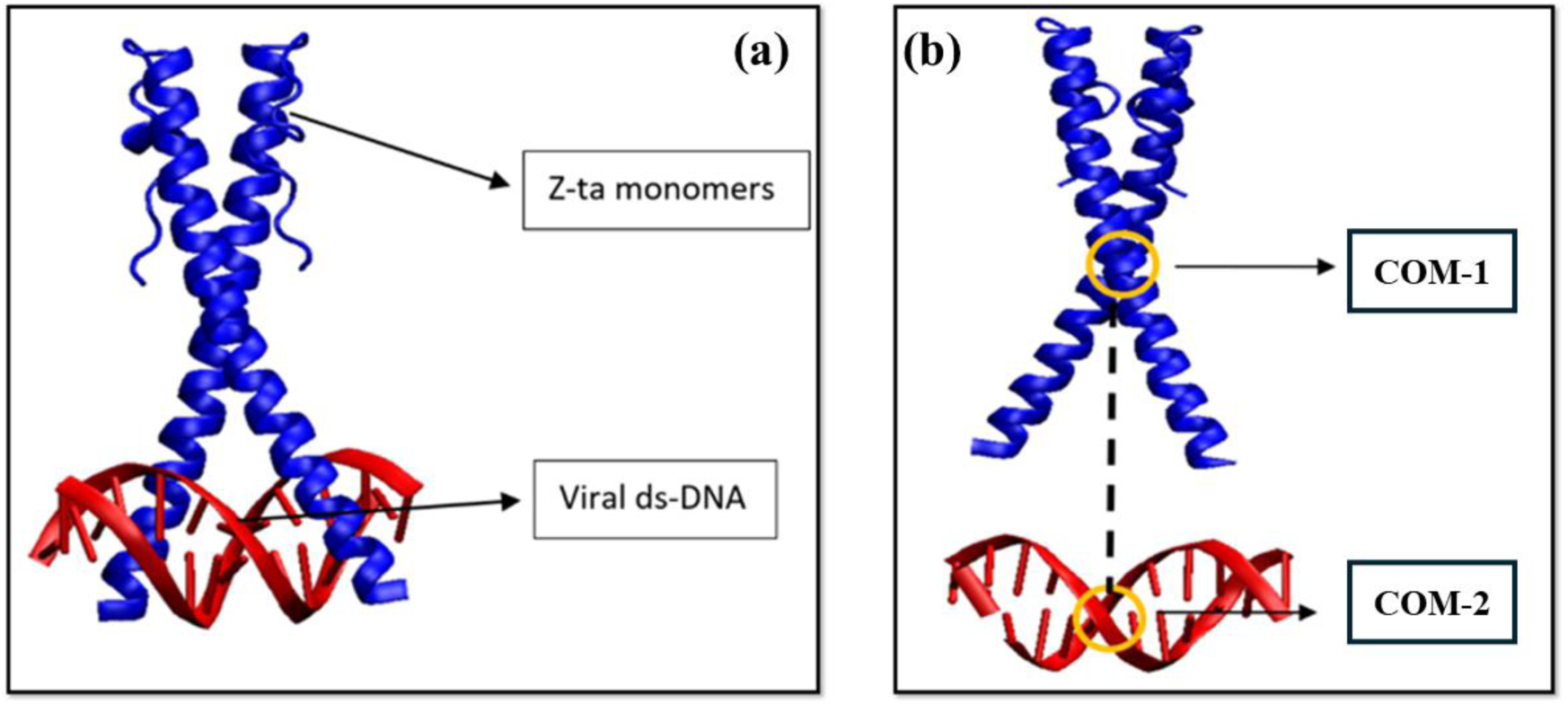
(a) It shows the modelled structure of the ZTA-DNA complex. The ZTA protein consists of two monomers (shown in blue colour), and a double stranded DNA (dsDNA) shown in red colour. (b) The centre of masses (COM) for two groups of atoms, protein (COM 1) and DNA (COM 2) respectively. The COM – COM distance between the protein and DNA groups is considered as the reaction coordinate to perform the umbrella sampling simulations.

**Table 1:** Details of three ZRE systems with ZTA protein studied in this work. The DNA nucleic base sequences are shown for three ZREs corresponding to core motif and core motif with flanking ends respectively.

| Protein | DNA sequences | Core motif sequences | Core motif with flanking ends sequences |
| --- | --- | --- | --- |
| ZTA | ZRE 1 | TGAGCCA | TTA-TGAGCCA-TTG |
|  | ZRE 2 | TGAGCGA | TTA-TGAGCGA-TTT |
|  | ZRE 3 | TTCGCGA | AAA-TTCGCGA-TGC |

### Molecular dynamics simulation

After the initial system setup, energy minimization was performed using the steepest descent algorithm^67^ with a force cutoff of 1000 kJ/mol/nm. The system’s energy was minimized by removing overlaps between the water molecules and solute in the system. Once the system was energy minimized, we performed NVT equilibration for 1 ns, followed by NPT equilibration for 1 ns. The system temperature was maintained at 310 K and the pressure at 1 bar. To equilibrate the system, we used the v-rescale (velocity rescaling) thermostat with a time constant of 0.1 ps and the Parinello-Rahman Barostat^68^ with isotropic scaling of box vectors. In our simulation, we employed the Leap-frog integrator^69^ to integrate Newton’s equations of motion. At every 500 steps (1 ps), the neighbour list was updated. The non-bonded interactions, including both electrostatics and van der Waals interactions, were calculated using the Verlet^70^ cutoff scheme with a cutoff distance of 1.4 nm. The LINCS^71^ algorithm was used to restrain the hydrogen bonds associated with hydrogen atoms, enabling us to use an integration timestep of 2 fs. To calculate the long range electrostatic interactions, we employed the Particle Mesh Ewald (PME)^72^ method with a cubic interpolation of order 4. Following equilibration, we performed a production run of 200 ns for all the systems. Details of the unbiased simulations are provided in Table 2. To calculate the equilibrium structural properties, the systems needed to be properly equilibrated. We calculated various equilibrium properties such as RMSD (root mean square deviation), RMSF (root mean square fluctuation), COM (centre of masses) – COM distance for the ZTA–dsDNA complex. We calculated hydrogen bonds (h-bonds) using GROMACS and interaction fingerprints using PROLIF (protein ligand interaction fingerprints) tool,^73^ which captures and quantifies various interactions between biomolecules. To calculate specific type of interactions, PROLIF uses a criterion along with a defined SAMRTS pattern. In supplementary information, we provided a detailed description of RMSD, RMSF, and the criteria for various interactions^73^.

**Table 2:** Details of box sizes, total number of atoms, water molecules, ions and simulation times (production run times) are provided.

| Protein | DNA sequences |  | Box sizes (nm<sup>3</sup>) | Total no. of atoms | No. of water molecules | No. of ions | Simulation time (ns) |
| --- | --- | --- | --- | --- | --- | --- | --- |
| ZTA | core motif | ZRE 1 | 12 x 8 x 12 | 113659 | 37025 | 144 | 200 ns |
|  |  | ZRE 2 | 12 x 8 x 12 | 113654 | 37020 | 144 | 200 ns |
|  |  | ZRE 3 | 12 x 8 x 12 | 113669 | 37025 | 144 | 200 ns |
|  | core motif with flanking ends | ZRE 1 | 12 x 8 x 12 | 113657 | 36880 | 144 | 200 ns |
|  |  | ZRE 2 | 12 x 8 x 12 | 113617 | 36880 | 144 | 200 ns |
|  |  | ZRE 3 | 12 x 8 x 12 | 113637 | 36887 | 144 | 200 ns |

### Umbrella sampling

To calculate the binding affinity between the ZTA protein and dsDNA, we performed enhanced sampling simulation using the umbrella sampling (US) technique. In this method, we chose a series of simulation windows along a pre-selected reaction coordinate, each representing a distinct spatial arrangement between the DNA and protein. Here, we selected one group of atoms in the ZTA protein and another group in the DNA. The centre of mass (COM) distance between these two groups was taken as the reaction coordinate (Figure 2(b)). A total of 55 bins (windows) were selected, with the minimum and maximum bins at 0 nm and 10 nm, respectively. Each system was equilibrated for 1 ns at the NPT ensemble, and then we performed umbrella sampling simulation for 2 ns successively. A force of 300 kJ/mol/nm^2^ was applied to restrain the system at each bin. Following a similar simulation protocol, we performed 10 independent US simulations for each ZTA–DNA system. Using WHAM^74^ (weighted histogram analysis method), we calculated the potential of mean force (PMF) along the reaction coordinate for each run and then averaged the PMF over the 10 independent runs, including error bars. Details of the systems for umbrella sampling simulations are provided in Table 3. To evaluate the average free energy, we first calculated the independent PMF using WHAM. We then extracted individual probabilities (p) using the relation *p* = exp (-F/*k_B_*T), where F is the free energy, *k_B_* is the Boltzmann constant, and T is the absolute temperature of the system. We then calculated the average free energy (F_avg_) = -*k_B_*T ln(*p_avg_*), where *p_avg_* is the averaged probability. The error bars were calculated using the formula = *k_B_*T(σ)/(sqrt(N).*p_avg_*), where σ is the standard deviation and N is the number of profiles.

**Table 3:**
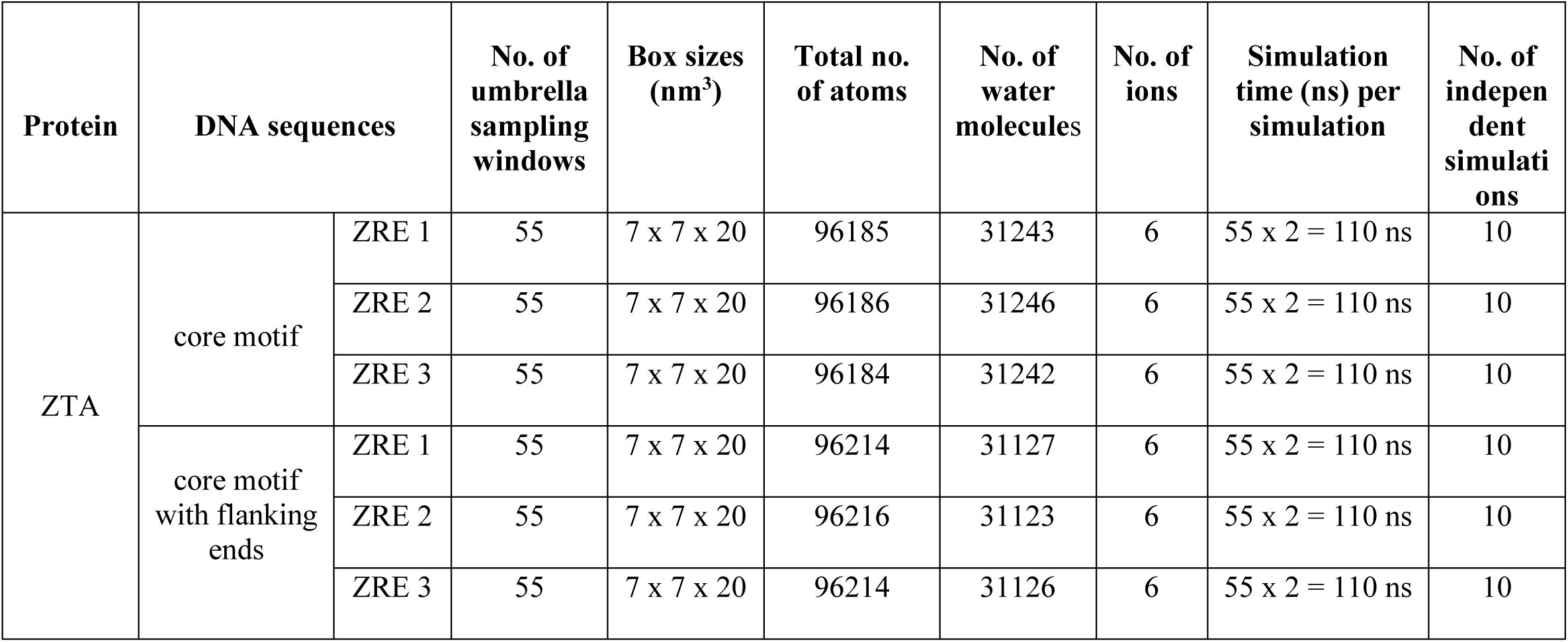
Details of umbrella sampling simulations for various systems with no of umbrella sampling windows, box sizes, total number of atoms, water molecules, ions, simulation times (production run times), number of independent simulations for each system are provided.

| Protein | DNA sequences |  | No. of umbrella sampling windows | Box sizes (nm<sup>3</sup>) | Total no. of atoms | No. of water molecules | No. of ions | Simulation time (ns) per simulation | No. of independent simulations |
| --- | --- | --- | --- | --- | --- | --- | --- | --- | --- |
| ZTA | core motif | ZRE 1 | 55 | 7 x 7 x 20 | 96185 | 31243 | 6 | 55 x 2 = 110 ns | 10 |
|  |  | ZRE 2 | 55 | 7 x 7 x 20 | 96186 | 31246 | 6 | 55 x 2 = 110 ns | 10 |
|  |  | ZRE 3 | 55 | 7 x 7 x 20 | 96184 | 31242 | 6 | 55 x 2 = 110 ns | 10 |
|  | core motif with flanking ends | ZRE 1 | 55 | 7 x 7 x 20 | 96214 | 31127 | 6 | 55 x 2 = 110 ns | 10 |
|  |  | ZRE 2 | 55 | 7 x 7 x 20 | 96216 | 31123 | 6 | 55 x 2 = 110 ns | 10 |
|  |  | ZRE 3 | 55 | 7 x 7 x 20 | 96214 | 31126 | 6 | 55 x 2 = 110 ns | 10 |

## RESULTS

### Structural stability of the ZTA TF – DNA complexes

To check the structural stability of the ZTA-DNA complexes for three different ZRE sequences - ZRE 1, ZRE 2 and ZRE 3 - we equilibrated each structure for 200 ns and calculated equilibrium properties such as RMSD (root mean square deviations), RMSF (root mean square fluctuations), and COM (centre of mass) distances, as shown in Figures 3-6. In Figure 3(a), for ZRE 1, the initial RMSD of the ZTA protein starts around 0.25 nm, and after 100 ns of simulation, it reaches approximately 0.45 nm and fluctuates within that range over the subsequent 200 ns of simulation time. For ZRE 2, it starts from 0.3 nm, initially increases to 0.7 nm within 100 ns of simulation, but then decreases and stabilizes around 0.45 – 0.5 nm. For ZRE 3, the initial RMSD value is around 0.4 nm; it fluctuates between 0.3 – 0.5 nm, and after around 100 ns of simulation, it saturates around 0.45 nm. From the RMSD plots for all three ZREs for the core motif cases, we observe that the RMSD for ZRE 3 is more stable, with less fluctuation over the 200 ns of simulation time. Thus, the ZRE 3 – ZTA TF complex shows better structural stability over the simulation time, as indicated by the RMSD analysis. For the core motif with flanking ends (Figure 3 (b)), except for ZRE 2, the RMSD values for ZRE 1 and ZRE 3 do not change much over the simulation time, remaining stable at around 0.3 nm after starting from approximately 0.25 nm. For the ZRE 2 system, the RMSD fluctuates between 0.25 – 0.5 nm until around 100 ns and then fluctuates around 0.35 nm over the remaining time. In Figure 4(a), the RMSD values of dsDNA for ZRE 1, ZRE 2 and ZRE 3 start from 0.2 nm and after 100 ns of simulations, saturate around an average RMSD of 0.35, fluctuating around 0.3 – 0.4 nm. Similarly, for the dsDNA of the core motif with flanking ends (Figure 4(b)), the RMSD starts around 0.2 nm, reaches 0.3 – 0.45 nm around 50 ns, and then fluctuates and saturates between 0.25 – 0.35 nm. Thus, for both cases and for all three ZRE systems, the change in RMSDs is within 0.2 nm, which is reasonably small. Hence, the dsDNA systems of the ZTA – DNA complex equilibrate over the course of the production run. In Figure 5, we show the COM distance between the ZTA protein and dsDNA COMs. In Figure 5(a), for the core motif ZRE cases, the COM distances initially fluctuate between 0.2-0.3 nm until 100- 125 ns. Later, the COM distances converge to around 2.8 – 3 nm over the simulation time. For the core motif with flanking ends (Figure 5(b)), the COM distances for all three ZREs saturate from the beginning, maintaining around 2.6 nm over the entire course of the simulation. The COM distance plot shows better structural stability for the core motif with flanking ends in comparison to the core motif cases alone. Figure 6 shows the root mean square fluctuation for the DNA chains for different ZRE systems. The RMSF plots show the fluctuations of each of these protein residues from their equilibrium conformations. As can be seen from Figure 6(a) and Figure 6(b), the overall RMSF fluctuation is lower, in the order of ∼ 0.1 nm, apart from the terminal nucleic acid bases for both the core motif and core motif with flanking ends, where the terminal ends show higher fluctuations. For ZRE 3 in the core motif with flanking ends, the structure is comparatively more stable than all other ZREs, highlighting the importance of the flanking ends in providing better stability to the core motif sequences. Thus, from the RMSD, RMSF, and COM plots, we have demonstrated the stability of the ZTA–dsDNA systems. Later, to quantitatively estimate the binding affinity between ZTA and dsDNA for various cases, we performed enhanced sampling (umbrella sampling simulations) and calculated free energies.

**Figure 3:**
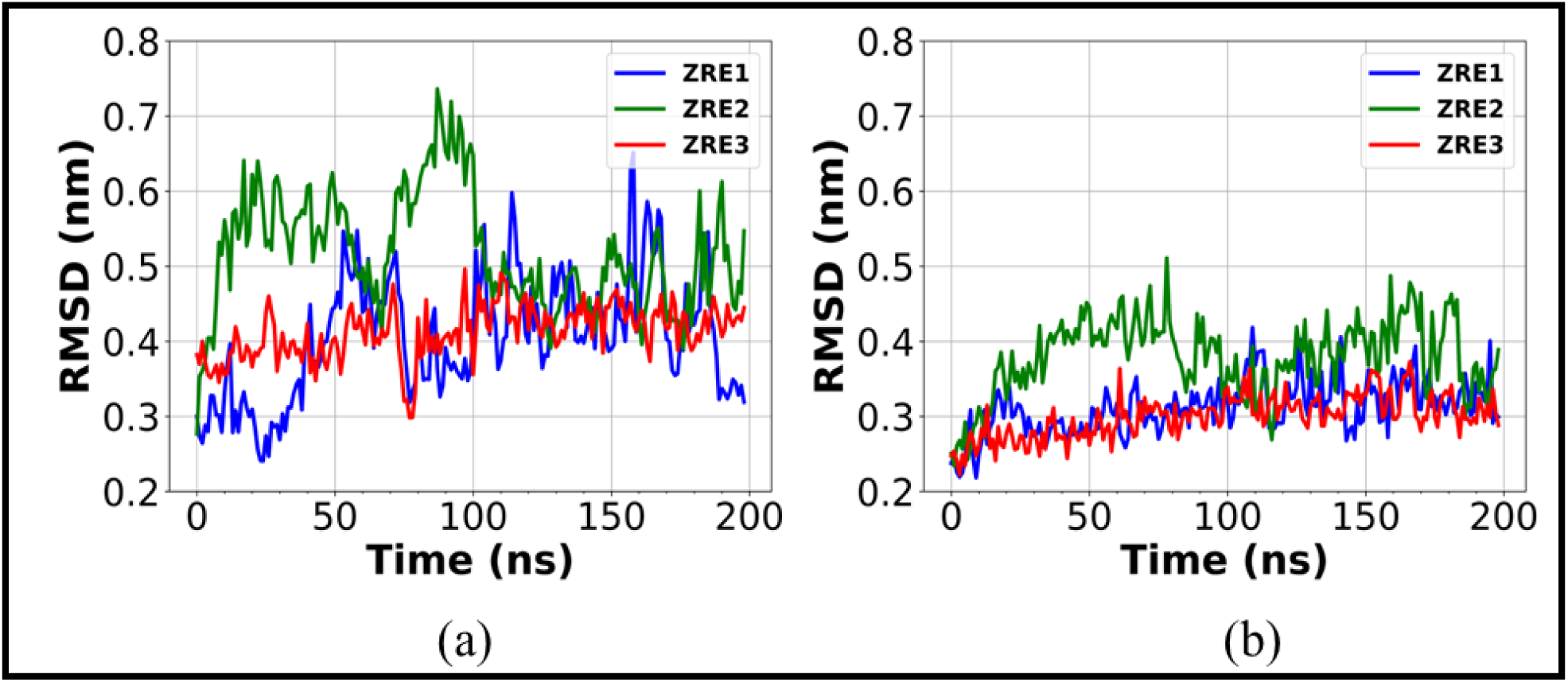
Figure shows RMSD (root mean square deviation) profiles of protein for both, (a) core-motif, and (b) core-motif with flanking ends, for three ZRE cases, ZRE 1 (blue), ZRE 2 (green), ZRE 3 (red) respectively. For each system, the RMSD is calculated from 200 ns unbiased MD simulation trajectories.

**Figure 4:**
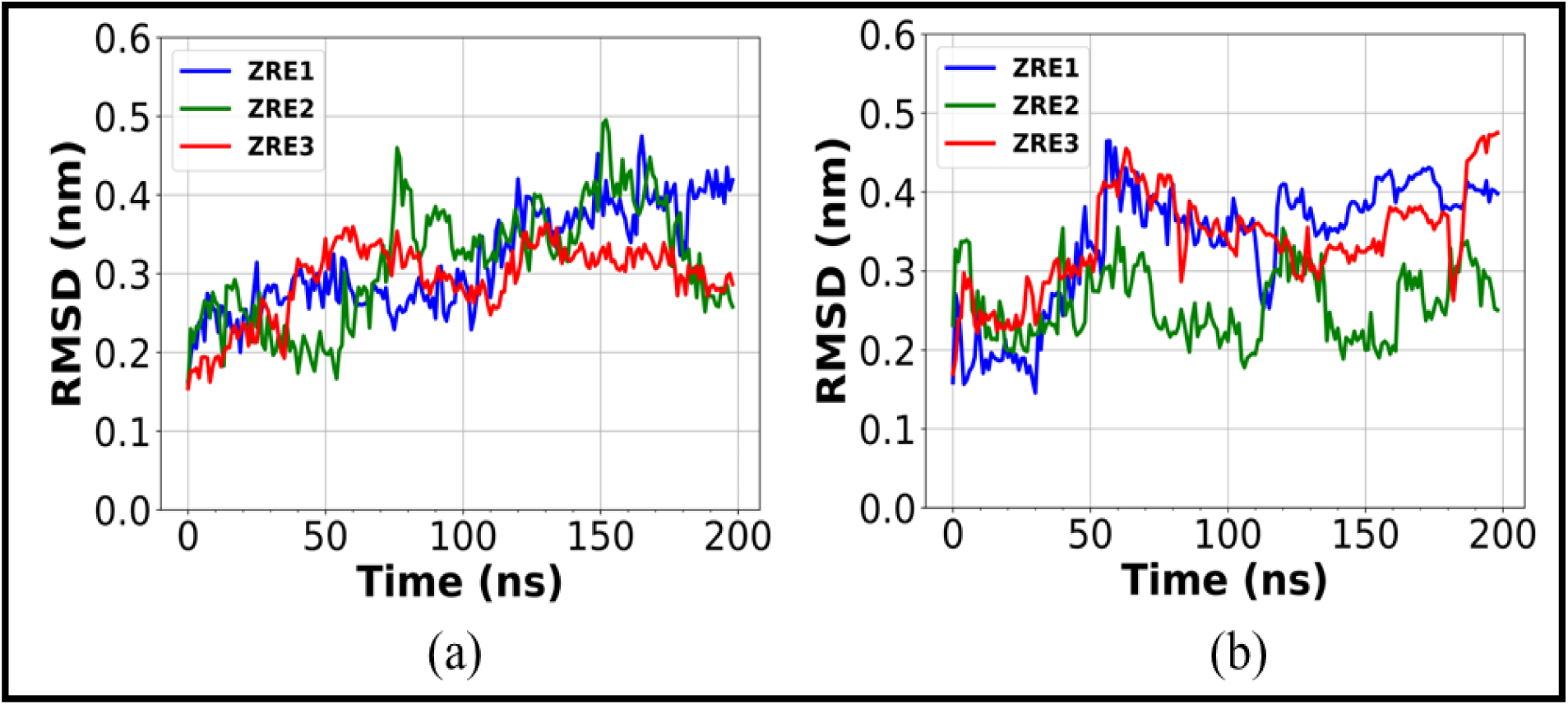
Figure shows RMSD (root mean square deviation) profiles of DNA for both, (a) the core motif, and (b) core motif with flanking ends for all three ZRE cases, ZRE 1 (blue), ZRE 2 (green), ZRE 3 (red) respectively. For each system, the RMSD is calculated from 200 ns unbiased MD simulation trajectories.

**Figure 5:**
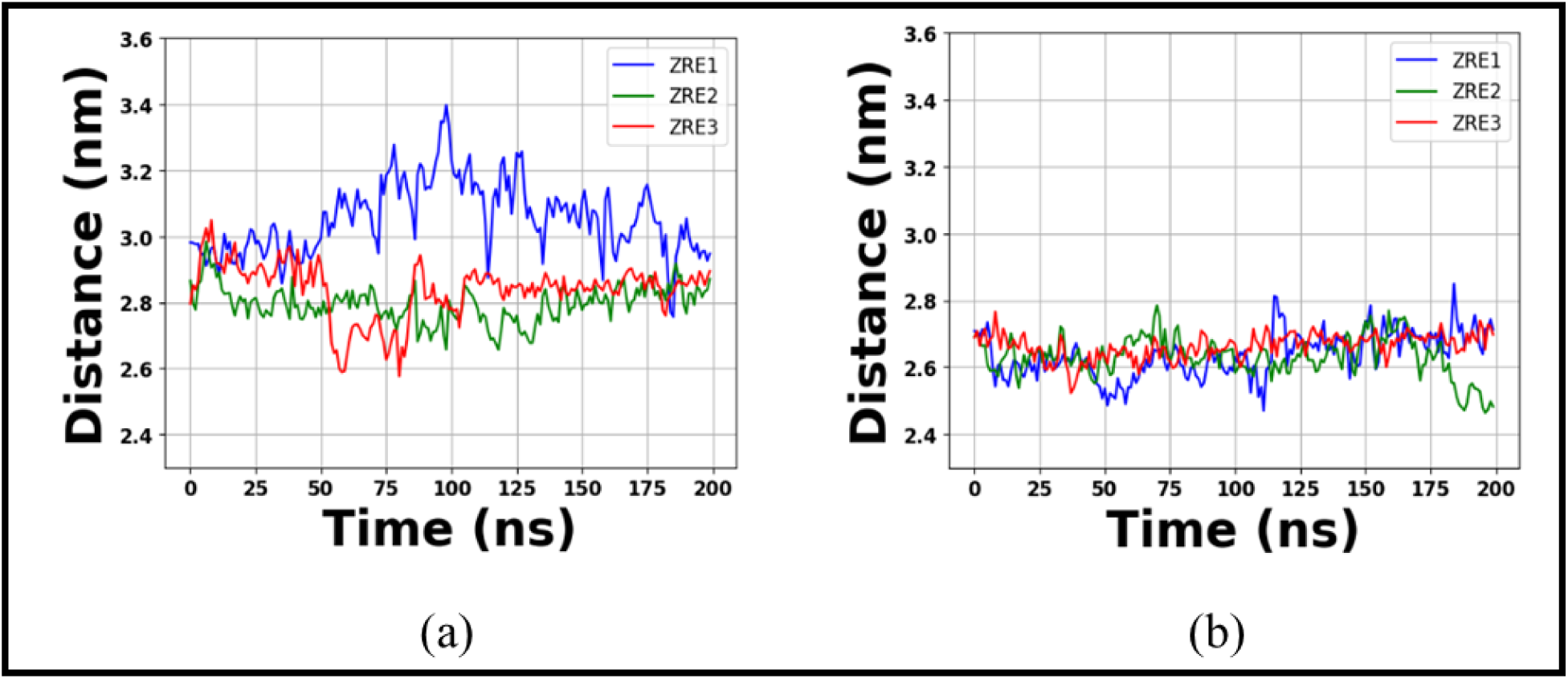
The centre of mass (COM) distance plot from the 200 ns unbiased simulation trajectories. The COM distance is calculated between protein and DNA COMs for both, the core motif and core motif with flanking ends for all three ZRE cases, ZRE 1 (blue), ZRE 2 (green), ZRE 3 (red) respectively. **(a)** COM distances for the core motifs. **(b)** COM distances for the core motif with flanking ends.

**Figure 6:**
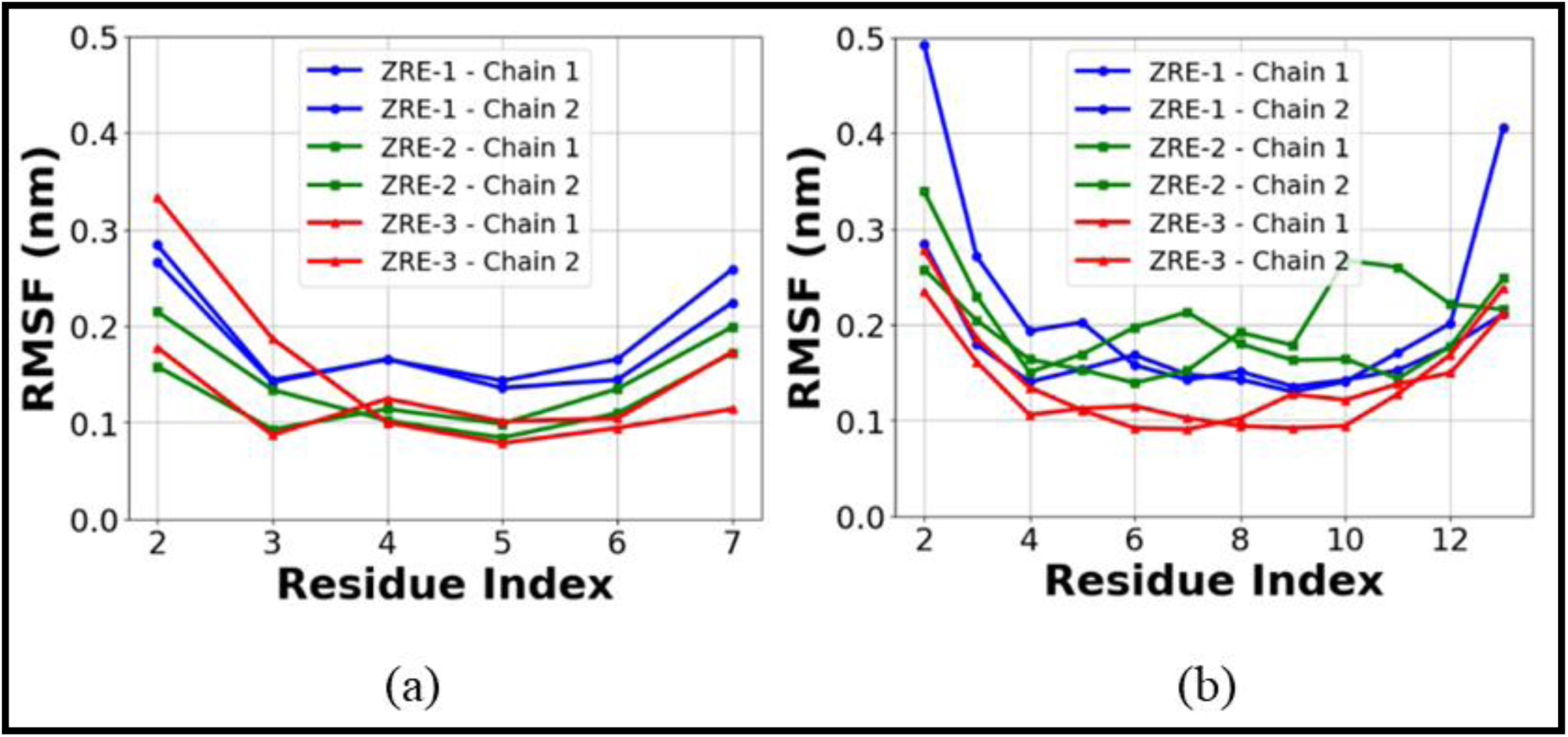
Root means square fluctuation (RMSF) plot of DNA chains for the (a) core motif and (b) core motif with flanking ends systems. For both the cases, different ZRE systems are shown in blue (ZRE 1), green (ZRE 2), and red (ZRE 3) colour respectively. For a specific ZRE system, we have denoted both the chains in same colour.

### Quantitative estimation of the binding affinity between ZTA TF and dsDNA

We report free energy surface (FES) plots with the centre of mass (COM) distance between the protein and DNA as the reaction coordinate, along with quantitative values of the free energy (Δ*G*) for ZTA TF and dsDNA systems from umbrella sampling simulations. Here, we show average FESs for the core motifs in Figure 7(a) and for the core motif with flanking ends in Figure 7(b). In Figure 7(a), we present the average FES profiles with error bars for different ZTA TF and dsDNA sequences. The FES profiles for ZRE 1 (blue), ZRE 2 (green), and ZRE 3 (red) are shown respectively for the core motif sequences, with Δ*G* values of approximately -33.47 ± 0.41 kcal/mol, -46.33 ± 0.36 kcal/mol, and -46.04 ± 0.25 kcal/mol (as shown in Table 4). In Figure 7(b), the FES profiles for ZRE 1 (blue), ZRE 2 (green) and ZRE 3 (red) are shown respectively for the core motif with flanking ends sequences, with Δ*G* values of approximately -67.87 ± 0.40 kcal/mol, -76.43 ± 0.50 kcal/mol, and -82.06 ± 0.34 kcal/mol (as shown in Table 4). Thus, in the absence of DNA flanking ends, we observe Δ*G* values in the range of approximately - 33 to -46 kcal/mol, whereas, with the addition of flanking ends to the core motif sequences, the Δ*G* values are in the range of approximately -67 to -82 kcal/mol. Therefore, the addition of flanking ends to the core motif sequences provides better stability to the dsDNA and ZTA TF complex, significantly lowering the free energy values for all ZRE systems. As shown in the FES profiles, the FES value for the ZRE 3 case is the lowest, indicating the best stability for the ZTA TF and dsDNA complex. In the insets of Figure 7(a-b), we show FES profiles at the global minima for the reaction coordinate values of 2 – 4 nm. In Figure 8, we present an instantaneous snapshot showing the bound and unbound states for the ZTA–dsDNA system. The bound and unbound states are illustrated with increasing COM – COM distance between the protein and DNA. We show the FES profile for the core motif with flanking ends systems for the three ZRE cases, and the snapshots represent the bound state corresponding to the global minima in the FES profile and the unbound state corresponding to the non-interacting region in the FES profile. The colours represent the electrostatic potential surface, with blue indicating positive charges and red indicating negative charges. This FES analysis shows that the interaction dynamics of ZTA-DNA complexes are significantly influenced by alterations in the end sequences of the dsDNA in conjunction with the core motif. In the supplementary information, we provide all the individual FES plots for all six cases, and for each system, we present all 10 independent FES profiles as shown in Figure S1(a-f).

**Figure 7:**
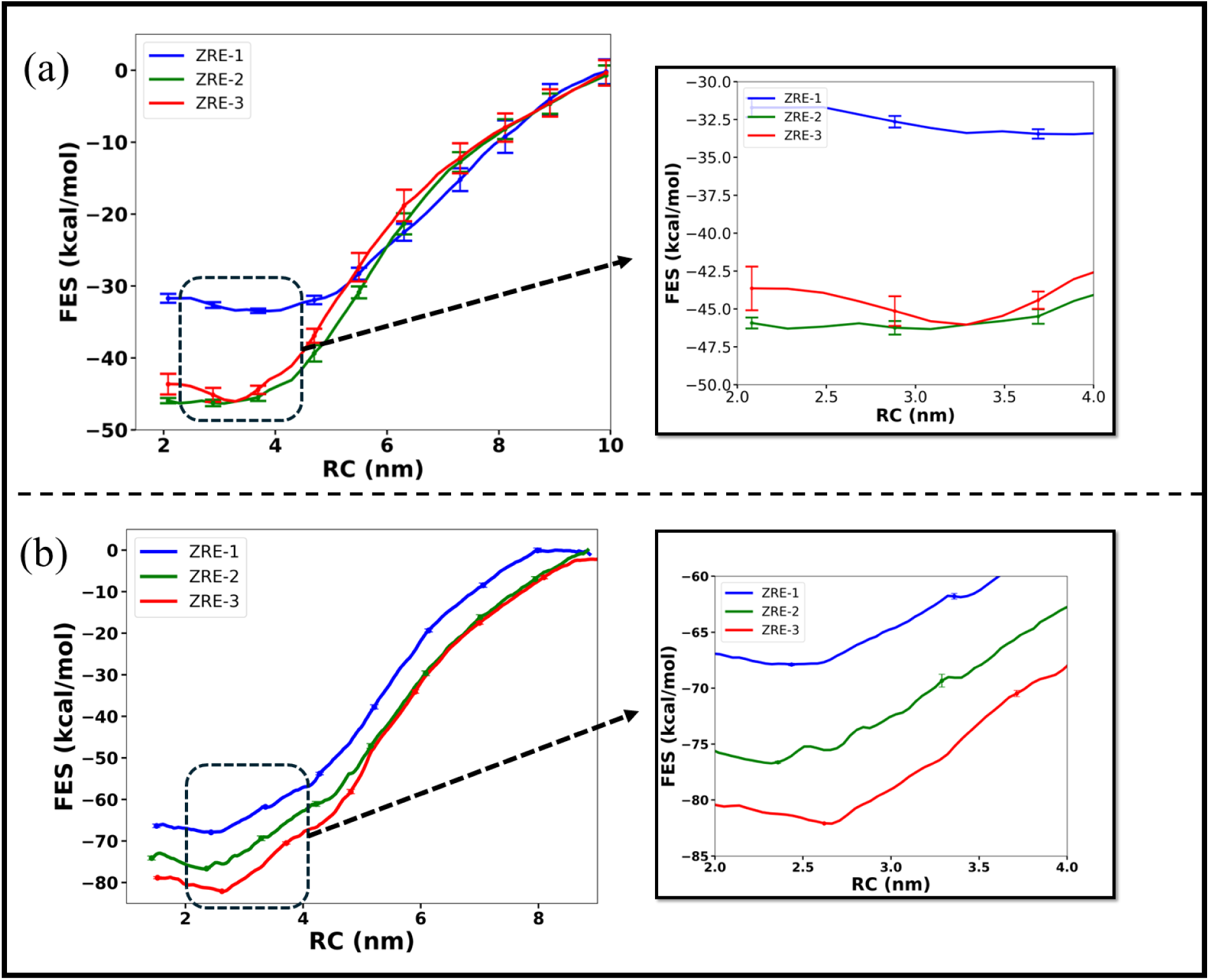
Average free energy surface (FES) plots with distance (COM – COM distance between protein and DNA) as reaction coordinate for the umbrella sampling simulations. Here we show average FESs for the (a) core motifs and (b) core motif with flanking ends. In both cases, different ZRE systems are shown in blue (ZRE 1), green (ZRE 2), and red (ZRE 3) respectively. FES plots are shown with error bars for all the cases. Inset shows zoomed in view of the FES at the global minima positions.

**Figure 8:**
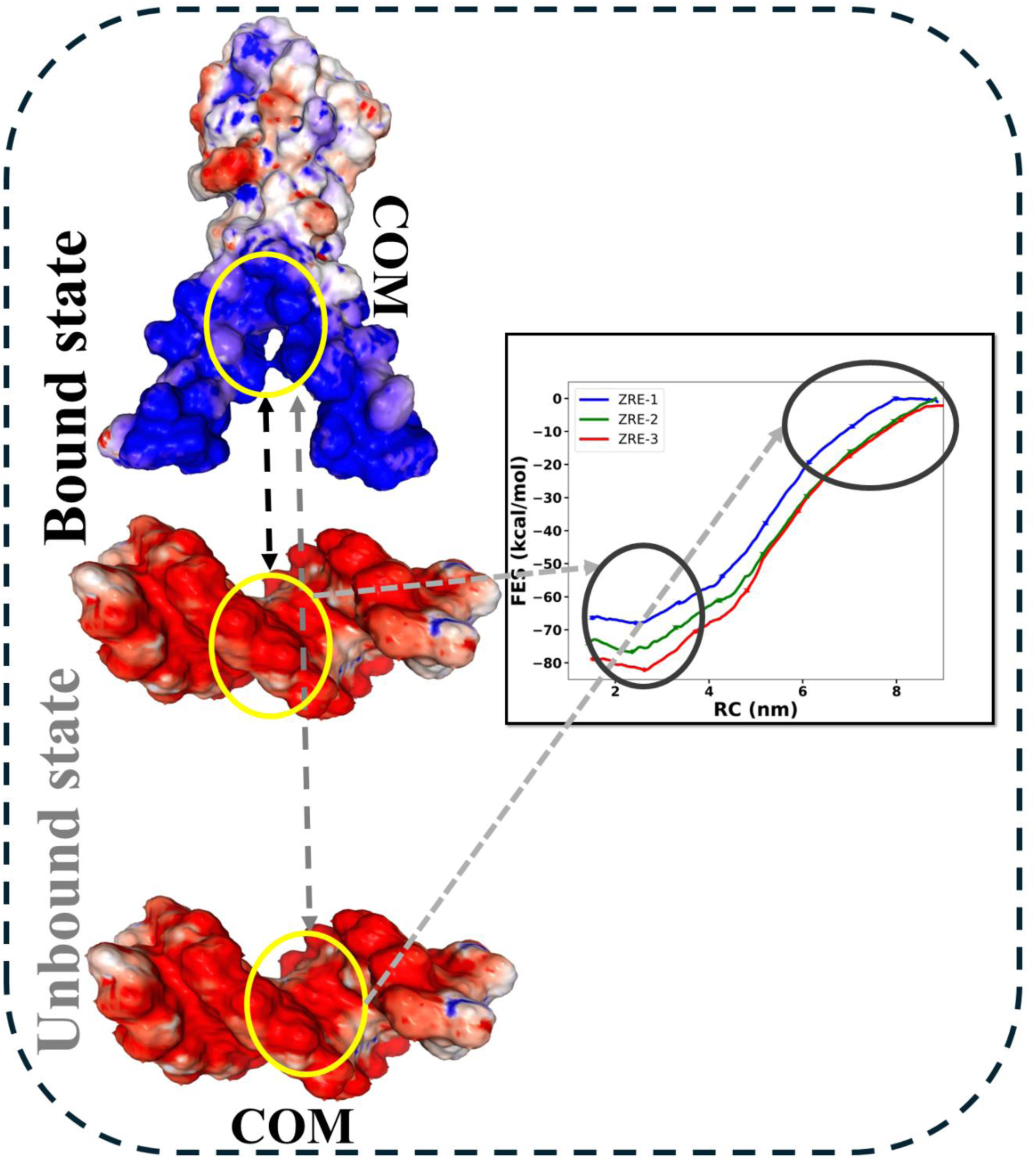
Instantaneous snapshot shows bound and unbound state for the ZTA – DNA system. The bound and unbound states are shown with increasing COM – COM distance between the protein and DNA. We show the FES profile for the core motif with flanking ends systems for three ZRE cases and the snapshots shown here represent the bound state corresponding to global minima in the FES profile and unbound state corresponding to the non-interacting region in the FES profile. The colours represent the electrostatic potential surface, with blue indicating positive and red indicating negative charges.

**Table 4:** Averaged free energy values with error bars for all systems calculated from umbrella sampling simulations.

| Protein | DNA sequences | | Free energy ± error bar (kcal/mol) |
| --- | --- | --- | --- |
| ZTA | core motif | ZRE 1 | $\sim -33.47 \pm 0.41$ |
| | | ZRE 2 | $\sim -46.33 \pm 0.36$ |
| | | ZRE 3 | $\sim -46.04 \pm 0.25$ |
| | core motif with flanking ends | ZRE 1 | $\sim -67.87 \pm 0.40$ |
| | | ZRE 2 | $\sim -76.43 \pm 0.50$ |
| | | ZRE 3 | $\sim -82.06 \pm 0.34$ |

### Key interactions providing structural stability to the ZTA TF and dsDNA complex

To delve into the basics of structural stability and the reasons behind the strong binding affinity between the ZTA TF and dsDNA complex, we calculated key interactions between ZTA and dsDNA components from unbiased MD simulation trajectories. To investigate the molecular - level interactions, we extracted hydrogen bond (h-bond) information using Gromacs^64^ and other interactions (anionic, van der Waals (VDW), and hydrophobic) using the ProLIF tool.

### Major protein residues and DNA nucleic bases forming interactions

Using ProLIF^75^, we calculated interaction fingerprints of the ZTA protein interacting with DNA sequences (ZRE 1, ZRE 2 and ZRE 3) for both the core motif and the core motif with flanking ends. Fingerprint analysis shows different types of interactions occurring over the simulation time between the ZTA protein and dsDNA. In the fingerprint calculations (shown in the supplementary information), we show all the interactions between ZTA and DNA. However, most of the interactions do not persist long enough and thereby die out, contributing ineffectively to the stability of the ZTA-dsDNA complex. Therefore, to identify the important interactions, we focussed on those occurring for more than 90% of the simulation time.

From the interaction fingerprint, we plotted bar charts to analyse the results. In Figure 9, we show the total counts of various interactions (anionic, van der Waals (VDW) and hydrophobic) occurring for more than 90% of the simulation time. We also highlighted the distribution of interactions occurring for 100% of the simulation time. As observed in Figure 9, for the core motif system, ZRE 2 shows the highest number of interactions persisting for 100% of the simulation time, followed by ZRE 3 and ZRE 1. For the core motif with flanking ends, ZRE 3 shows the highest number of persistent interactions for 100 % of the time, with ZRE 2 and ZRE 1 following. Overall, for the core motif case, the total number of interactions follows the trend: ZRE 2 > ZRE 1 > ZRE 3, and for the core motif with flanking ends, ZRE 3 > ZRE 2 > ZRE 1 for the 90% time. For the 100% time, in the core motif case, we observe few interactions at 100% for the ZRE 2 and ZRE 3 cases, whereas for the core motif with flanking ends, we observe 100% interaction only for ZRE 3 but not for ZRE 1 and ZRE 2. The maximum number of total interactions above 90% is 14 for the core motif case and 13 for the core motif with flanking ends. From Figure 9, we can conclude that for the core motif cases (ZRE 1, 2, and 3), there are specific trends; however, for the core motif with flanking ends, we observe that ZRE 3 has the highest number of interactions (90 % as well as 100 %), providing better stability.

**Figure 9:**
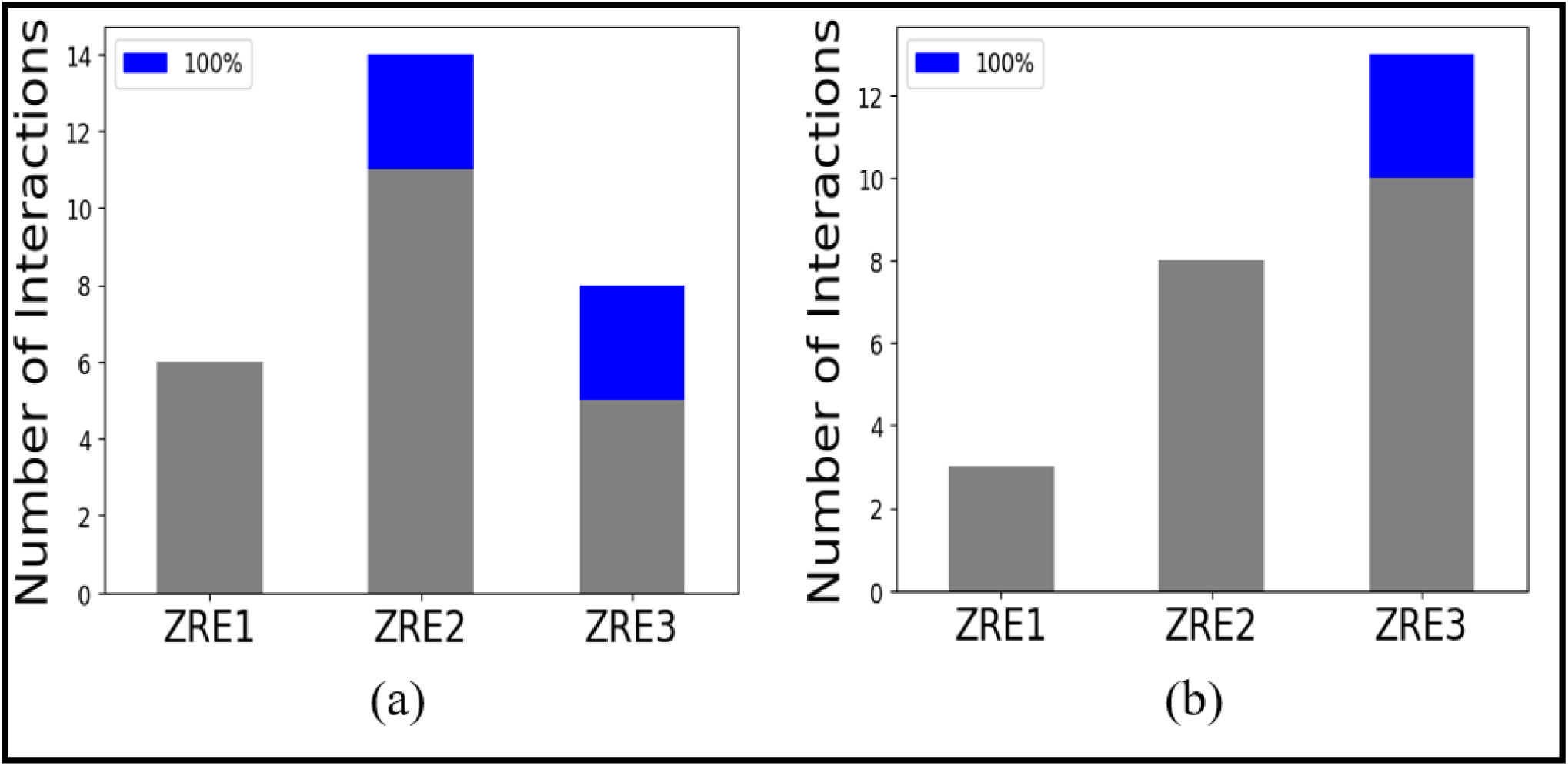
Overall numbers of different types of dominant interactions (anionic, van der Waals (VDW), hydrogen bond, and hydrophobic) between ZTA protein and DNA for ZRE 1, ZRE 2 and ZRE 3 systems respectively. The interactions have been calculated from unbiased MD simulation trajectories. Interactions which stay for more than 90 % of the simulation’s times are accounted here. Also, the one which stays for 100 % of simulation time are highlighted in blue colour on top of grey (numbers of interactions having more than 90% simulation times). Figure (a) for core motif systems, and (b) core motif with flanking ends, for three ZRE cases respectively.

In Figure 10, we explored the quantitative estimation of these different types of interactions (anionic (blue), van der Waals (VDW) (orange), hydrogen bond (green), and hydrophobic (red)) between ZTA and DNA systems calculated from unbiased MD simulation trajectories for (a) the core motif and (b) the core motif with flanking ends, for the three ZRE cases respectively. The interactions present for more than 90% of the total simulation times were counted here. As observed in Figure 9, similar trends are followed here. The van der Waals interaction is the most dominant interaction across all cases. For ZRE 2 core motif, there are 7 van der Waals interactions, which is approximately 50% of its total counts (7 out of 14 total interactions). Hydrogen bonds and anionic interactions are also the next dominant interactions (20%), while hydrophobic interaction is almost negligible. For ZRE 3 core motif, out of a total of 8 interactions, 5 are van der Waals (∼ 60%), and anionic and hydrogen bond acceptors are ∼ 20–25% or less. In all cases, hydrophobic interaction doesn’t play a significant role. For ZRE 1 core motif, van der Waals, anionic, and hydrogen bond acceptors are each at 33% . For the core motif with flanking ends, like total counts, van der Waals interaction follows the trend of ZRE 3 (40%) > ZRE 2 (35%) > ZRE 1 (70%). For the ZRE 3 case, anionic and hydrogen bond acceptors are the next dominant interactions, similar for ZRE 2 as well, whereas for the ZRE 1 case, anionic interaction is present 30% only. Therefore, van der Waals interaction is the most dominant and major interaction, with >50% for all cases, which plays a key role in providing structural stability to the ZTA–dsDNA systems for both the core motif and core motif with flanking ends. In the supplementary information (Figure S2-S7), we provided fingerprint analysis for all these systems separately.

**Figure 10:**
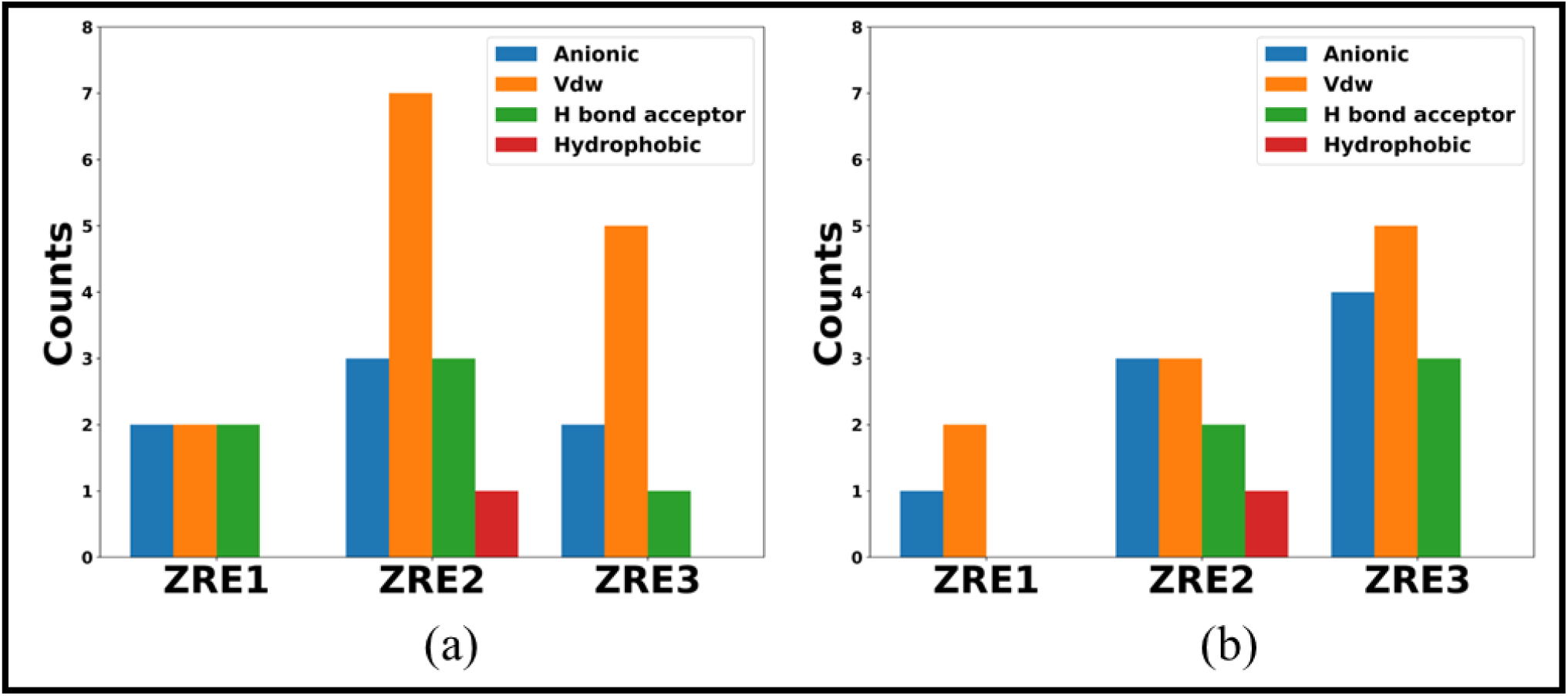
Different types of interactions (anionic (blue), van der Waals (VDW) (orange), hydrogen bond (green), and hydrophobic (red)) between ZTA – DNA systems calculated from unbiased MD simulation trajectories for (a) core motif and (b) core motif with flanking ends, for three ZRE cases. The interactions present over 90% of the total simulation times were counted here.

To get a better picture of the specific ZTA protein residues and DNA nucleic bases that participate in the dominant interactions, we performed a detailed analysis. In Figure 11, we calculated the protein residues that form interactions with their DNA counterparts. As shown in Figure 11, we observe that Arginine is the main dominant protein residue, participating significantly in all the different types of interactions for over 90% of the total simulation time. Although there are interacting residues like alanine, asparagine, serine in the ZRE 3 core motif, and cysteine and serine in the ZRE 3 core motif with flanking ends, for ZRE 2 and ZRE 1, we do not observe any residues other than Arginine existing for more than 90 % of the total simulation time. Clearly, arginine plays the most important role in forming different types of interactions for all these cases. Similarly, to explore the dominant nucleic bases in DNA, we calculated and analysed all the interactions between ZTA TF and dsDNA, counting the number of interactions that persist for more than 90% of the total simulation time. In Figure 12, we show the counts of nucleic bases A, T, G, and C that participate in the interaction with ZTA TF. As observed, except for ZRE 1 (core motif), in the other cases, any two, three, or all four bases (A, T, G, C) form interactions with protein residues. For the ZRE 1 core motif, only the nucleic base T participates in the interaction. For ZRE 2 (core motif), A and T form major interactions; for ZRE 3 (core motif), C and G interact; for ZRE 1 (core motif with flanking ends), the nucleic base G participates in interaction; for ZRE 2 (core motif with flanking ends), the nucleic base T, G and C interact; and for ZRE 3 (core motif with flanking ends), the nucleic bases G and A interact with ZTA TF. From this nucleic acid-level detailed analysis, we observe that nucleic acid G plays a dominant role in interactions for the ZRE 3 core motif with flanking ends system. Our free energy values and number of interactions show very good agreement for the ZRE 3 core motif with flanking ends (lowest free energy and highest number of interactions). Additionally, the presence of a higher number of G interactions possibly contributes to this highest stability, along with the highest number of total interactions. Later, we performed nucleic acid-level analysis to show how the presence of specific nucleic acids bases or combination of bases provide enhanced stability to the ZTA–dsDNA system. In Figure 13, we provided instantaneous snapshots of the ZTA–DNA complex for the ZRE 3 (core motif with flanking ends) to show the microscopic level picture at the all-atom level. The major protein residues interacting with the DNA at the ZTA binding domain are highlighted in blue, and the DNA nucleic bases are highlighted in red. In Figure 13 (a), we mainly shown arginine residues in the protein (dominant residues forming the interaction) interacting with the nucleic bases G or C. In Figure 13 (b), we show an instantaneous snapshot of the microscopic position of the 8 arginine residues (Arg 179, Arg 183, Arg 187, and Arg 190 in both ZTA monomers) which are the most dominant protein residues forming various interactions with the DNA strands.

**Figure 11:**
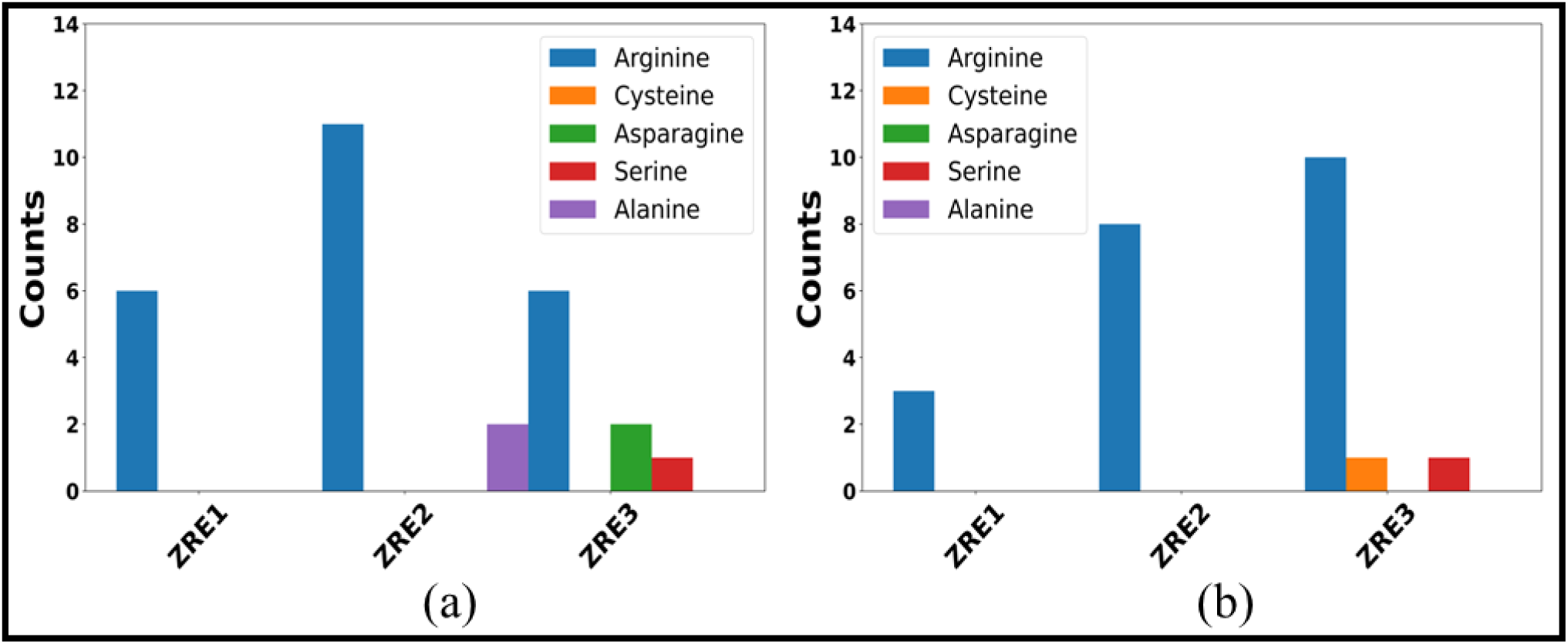
The plot shows amino acid residues from the ZTA protein which form various dominant interactions with DNA. The numbers of different amino acid residues in the y-axis are calculated from unbiased MD simulation trajectories and here we provided only those residues of the ZTA protein which forms interactions with DNA and stays for more than 90% of the total simulation times. In figure **(a)** we show core motifs, and (b) core motif with flanking ends for ZRE 1, ZRE 2 and ZRE 3 cases. Dominantly interacting amino acid residues are presented in blue (Arginine), orange (Cysteine), green (Asparagine), red (Serine), and violet (Alanine) respectively.

**Figure 12:**
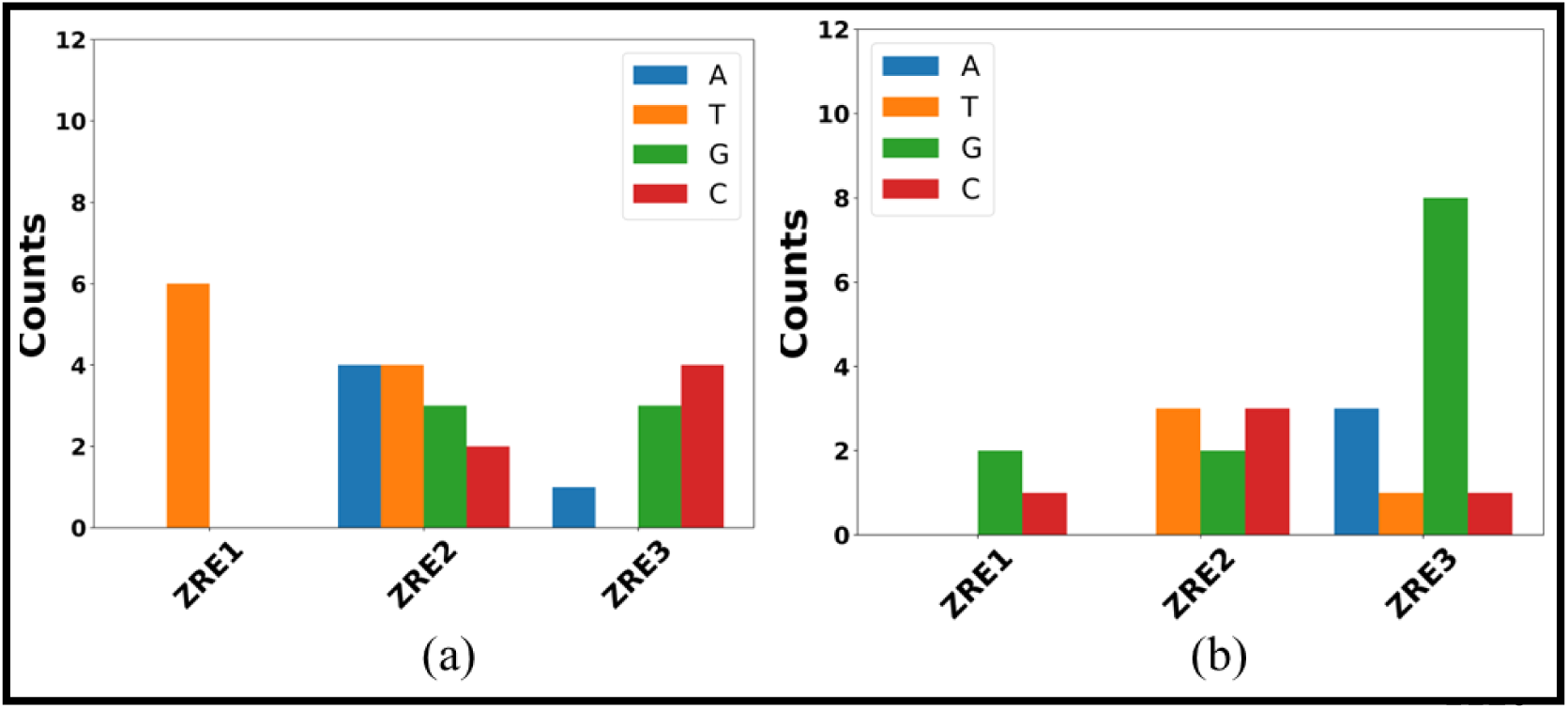
The plot shows count of four different types of DNA nucleic bases involved in interactions with ZTA protein. The numbers of different nucleic bases in the y-axis are calculated from unbiased MD simulation trajectories and only those nucleic bases are counted from the DNA which forms interactions with ZTA protein and stays for more than 90% of the total simulation times. In figure **(a)** we show core motifs, and (b) core motif with flanking ends for ZRE 1, ZRE 2 and ZRE 3 cases. The nucleic acid bases are presented in blue (adenine (A)), orange (thymine (T)), green (guanine (G)), and red (cytosine (C)) respectively.

**Figure 13:**
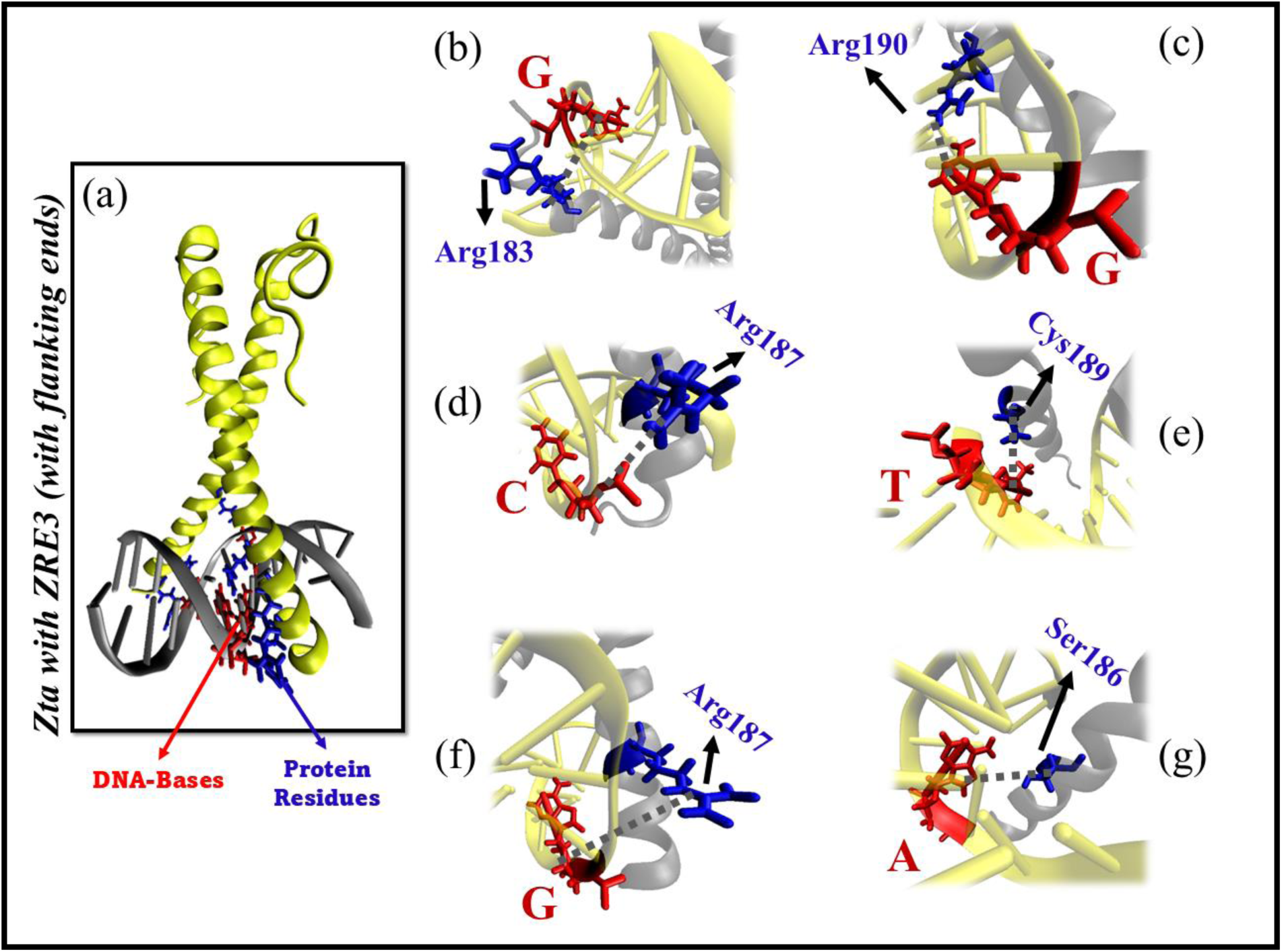
(a) Instantaneous snapshot shows DNA (grey colour) and ZTA protein (yellow) system for core motif with flanking ends for the ZRE 3 case. The protein residues interacting with DNA at the ZTA binding domain are highlighted in blue colour and DNA bases in red. Here we presented dominant interactions that occurs more than 95% of the simulation time. (b)Arg183 has 3 interactions at 100% of occurrence with G from DNA. (c, d, e, f) R190, R187, C189, R187 residues having interaction with DNA bases G, C, T, G respectively more than 95% of occurrence.

**Figure 13b:**
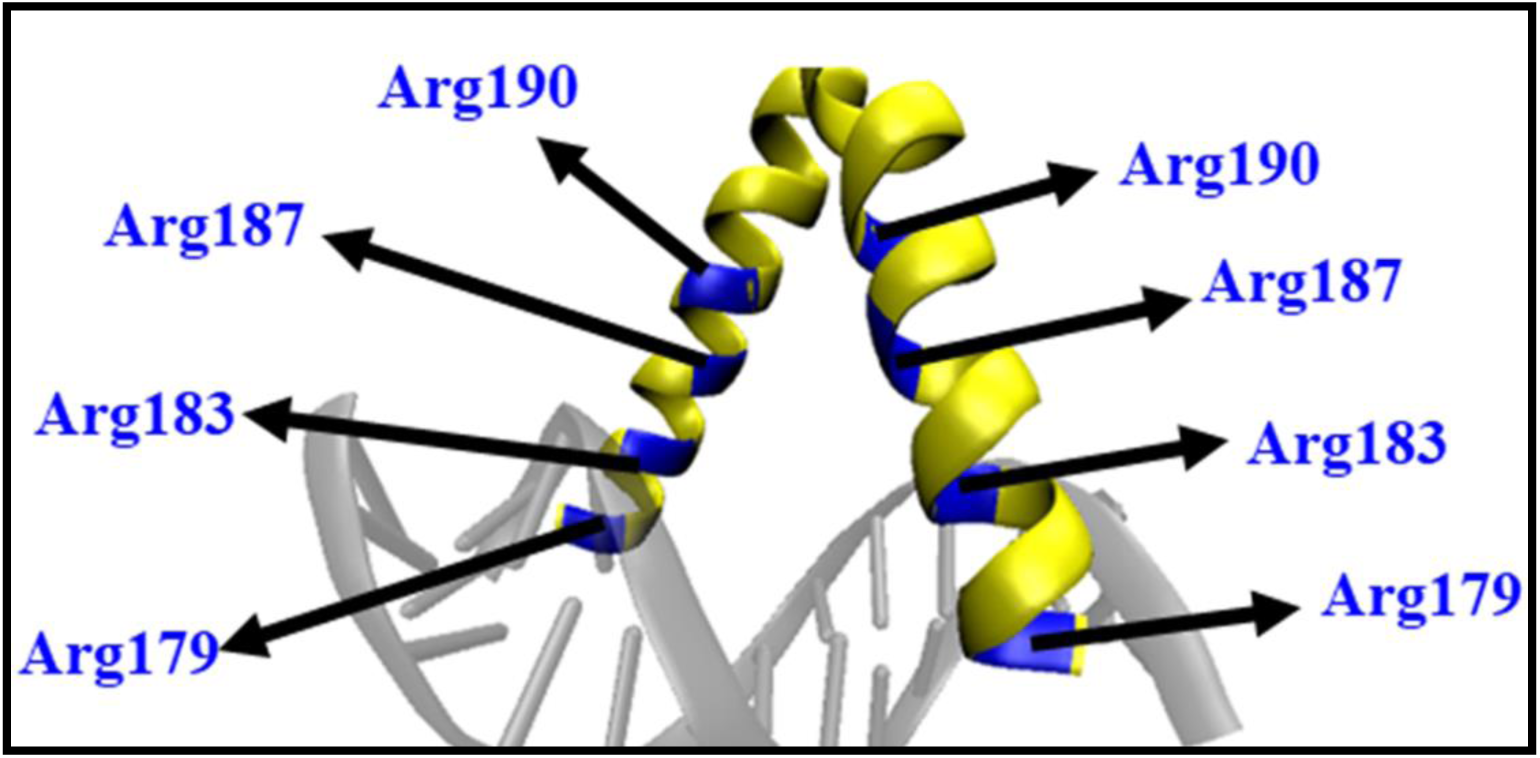
(b) Dominant ZTA residues Arginine (Arg) at the DNA binding domain from ZTA protein.

To explore the effect of nucleic base modification on the structural stability and binding affinity of the ZTA–dsDNA complex, we first explicitly counted the DNA nucleic bases predominantly involved in the interaction with the ZTA protein as shown in Figure 12. As previously reported by Hong et al.,^32^ the presence of C and G nucleic acid bases provides better stability to the ZTA–dsDNA complex. We calculated the number of C and G bases present in the core motif and core motif with flanking ends in all three ZRE systems respectively to check for any correlation between the free energy values and the number of C/G bases. In Figure 14, we calculated the number of dominant C and G bases in the dsDNA and schematically represented them. The cytosine (C) and guanine (G) nucleic acid bases from the dsDNA participate predominantly in the interaction with the ZTA protein. In Figure 14 (a), for ZRE 1, there are no C and/or G bases presents; for ZRE 2, there are two G and one C base; for ZRE 3, there are two G and two C bases. For the core motif with flanking ends, ZRE 1 has two C bases; ZRE 2 has two C and one G base; and for ZRE 3, there are two C and two G bases. Therefore, for both the core motif and core motif with flanking ends, the numbers of CG bases/pairs follow the pattern: ZRE 3 > ZRE 2 > ZRE 1. For the core motif case, we observe that for ZRE 2 and ZRE 3, both have two G bases, whereas the number of C bases is one (ZRE 2) and two (ZRE 3) respectively. For the core motif with flanking ends, for all three ZREs, there are two C bases, while the number of G bases is zero (ZRE 1), one (ZRE 2), and two (ZRE 3) respectively. Our calculated ΔG values for the core motif sequences are ZRE 1 ∼ -33.47 ± 0.41 kcal/mol, ZRE 2 ∼ -46.33 ± 0.36 kcal/mol, ZRE 3∼ -46.04 ± 0.25 kcal/mol (as shown in Table 4), following a similar trend considering the number of CG bases. Even for the core motif with flanking ends, the Δ*G* values are ZRE 1 ∼ -67.87 ± 0.40 kcal/mol, ZRE 2 ∼ -76.43 ± 0.50 kcal/mol, ZRE 3∼ - 82.06 ± 0.34 kcal/mol (as shown in Table 4), again following a similar pattern in CG base numbering: 2, 3, and 4 respectively. However, we can attribute the difference in Δ*G* in all the ZRE systems for the core motif with flanking ends to the presence of two C bases, which are not observed otherwise in the core motif cases. In both cases, ZRE 3 has a lower ΔG, having two CG pairs in each. Hence, although there are two CG pairs in ZRE 3, the presence of flanking ends provides better stability to the ZTA–dsDNA complex.

**Figure 14:**
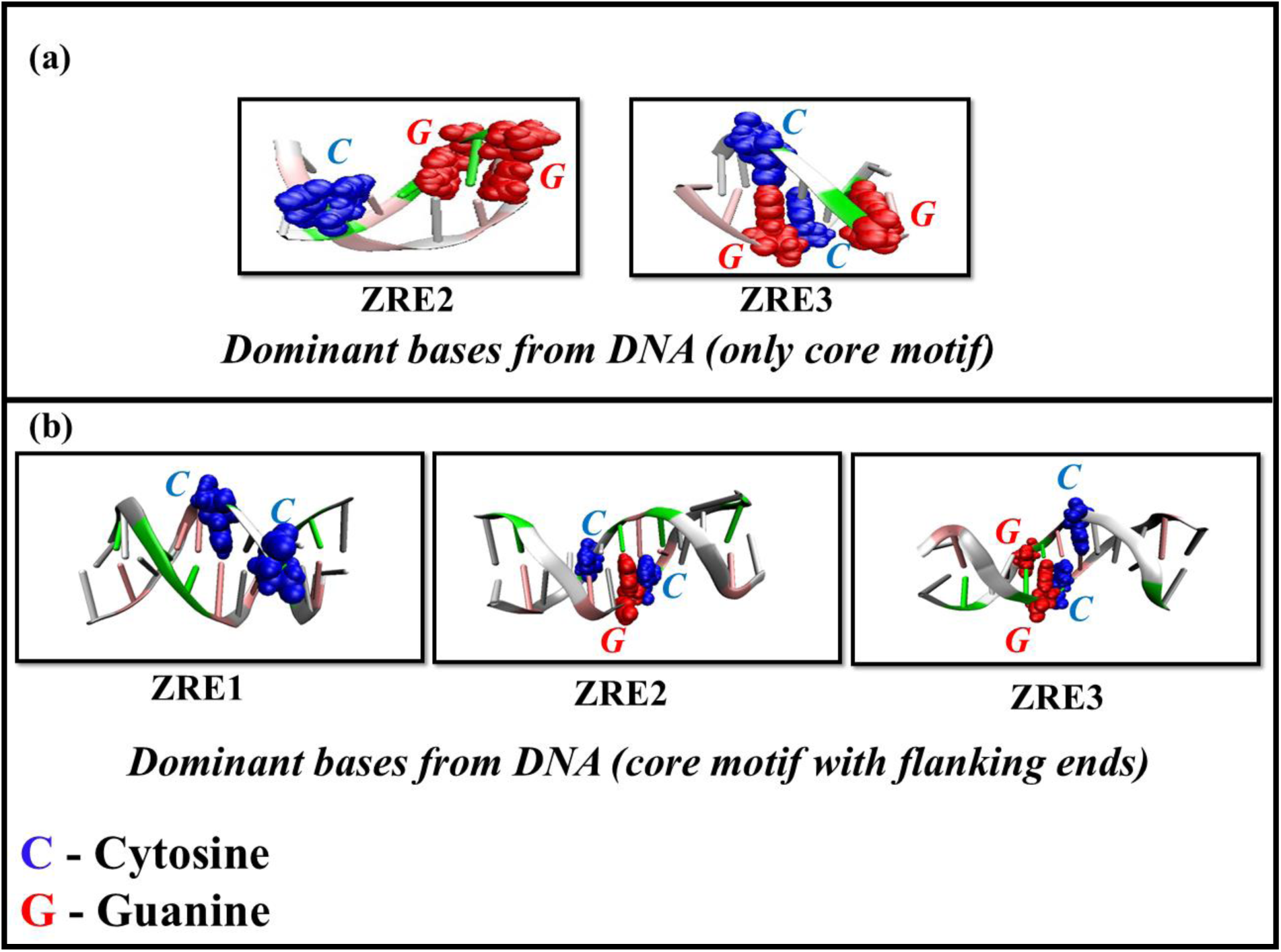
Cytosine (C) and guanine (G) nucleic acid bases from the dsDNA which takes part majorly in the interaction with ZTA protein. (a) Core motif case for ZRE2 and ZRE3. (b) Core motif with flanking ends for ZRE1, ZRE2 and ZRE3 respectively. The C bases are shown in blue colour and G in red colour.

### H-bond analysis between ZTA and DNA complex

Exploring full interaction details is paramount to elucidating the stability of ZTA–dsDNA complex systems. Earlier, in Figure 9 and Figure 10, we presented the total interaction counts and the counts of various types of interactions (anionic, cationic, van der Waals (VDW), hydrogen bonds, and hydrophobic interactions). We further calculated hydrogen bonds between the ZTA TF and dsDNA complex to investigate how hydrogen bonding varies between the core motif and the core motif with flanking ends across the three ZRE systems. Using unbiased MD simulation trajectories, we calculated the probability of hydrogen bonds between the ZTA protein and dsDNA strands. The probability of hydrogen bonds, along with the number of hydrogen bonds, is shown in Figure 15: (a) represents the core motif cases, and (b) represents the core motif with flanking ends. For ZRE 1 (core motif), the number of hydrogen bonds is approximately 12 with a probability of 0.13; for ZRE 2 (core motif), the number is around 18 with a probability of 0.14; and for ZRE 3 (core motif), the number is approximately 18 with a probability of 0.12. Thus, the hydrogen bond count follows the trend ZRE 1 (∼ 12) < ZRE 2 (∼18) ≤ ZRE 3 (∼18) for the core motif case, similar to the trend observed in the free energy values (ΔG) for the core motif sequences: ZRE 1 ∼ -33.47 ± 0.41 kcal/mol, ZRE 2 ∼ -46.33 ± 0.36 kcal/mol, and ZRE 3 ∼ -46.04 ± 0.25 kcal/mol (as shown in Table 4). For the core motif with flanking ends, Figure 15 (b) shows that ZRE 1 has approximately 19 hydrogen bonds with a probability of 0.13; ZRE 2 has around 18 with a probability of 0.13; and ZRE 3 has approximately 22 with a probability of 0.16. Here, the hydrogen bond count follows the trend ZRE 1 (∼ 19) ≤ ZRE 2 (∼18) < ZRE 3 (∼22) for the core motif with flanking ends. The trend in free energy values (ΔG) for the core motif with flanking ends is ZRE 1 ∼ - 67.87 ± 0.40 kcal/mol, ZRE 2 ∼ -76.43 ± 0.50 kcal/mol, and ZRE 3∼ -82.06 ± 0.34 kcal/mol (as shown in Table 4). For ZRE 1 and ZRE 2, despite a free energy difference of ∼ 10 kcal/mol, the hydrogen bond count remains the same. This indicates that other interactions between ZTA and dsDNA contribute to the difference between ZRE 1 and ZRE 2 in the core motif with flanking ends. However, ZRE 3 shows a notably higher hydrogen bond count (∼ 22) than all other ZRE cases, whether core motif or core motif with flanking ends, and also has the lowest ΔG value. Therefore, the hydrogen bond count shows a good correlation with free energy values in both cases, across different ZRE systems. This suggest that hydrogen bonding is one of the critical interactions contributing to the stability of the ZTA–dsDNA complex.

**Figure 15:**
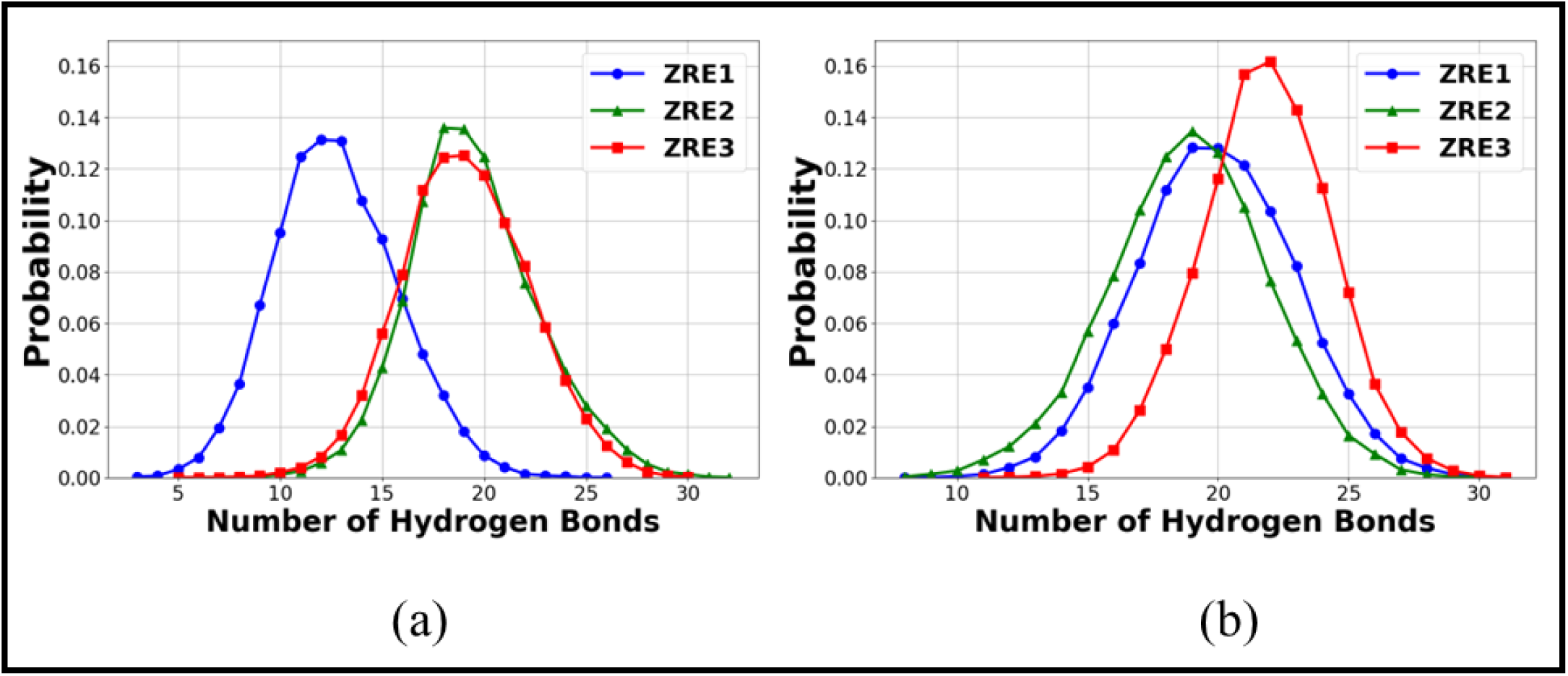
Probability of hydrogen bonds between ZTA protein and DNA nucleic bases. Figure (a) core motif, and (b) core motif with flanking ends, for ZRE 1 (blue), ZRE 2 (green) and ZRE 3 (red) respectively. The hydrogen bonds are calculated from the unbiased MD simulation trajectories.

### Distance analysis between ZTA monomer ends

To understand the effect of different core motif sequences for the three ZRE systems, we further studied the end-to-end distance between the two ZTA monomers, as shown in Figure 16. We calculated the distance between the two *α*-carbons of residue K176 from the two monomers and examined its time evolution over 200 ns unbiased MD simulation trajectories. In Figure 16, we show the distance calculation for the core motif (blue) and core motif with flanking ends (red), respectively, for (a) ZRE 1, (b) ZRE 2, and (c) ZRE 3 systems. The distances were as follows: for the core motif, ZRE 1 ∼ 22.43 (4.25) Å, ZRE 2 ∼ 32.94 (3.53) Å, and ZRE 3 ∼ 22.93 (2.96) Å; and for the core motif with flanking ends, ZRE 1 ∼ 26.79 (3.17) Å, ZRE 2 ∼ 26.79 (3.17) Å, and ZRE 3 ∼ 26.45(2.25) Å. Therefore, we observe that for the core motif cases, interchain distances do not change much over time, whereas for the core motif with flanking ends, the interchain distance decreases over time. Thus, the addition of flanking ends likely makes the ZTA chains more rigid.

**Figure 16:**
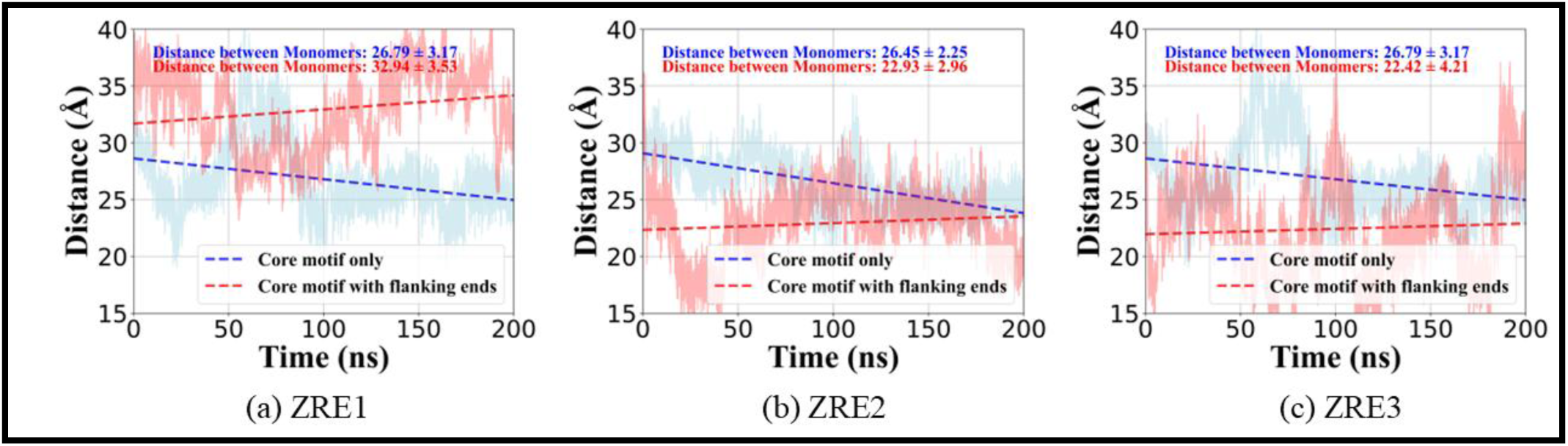
Distance vs time plot for core motif and core motif with flanking ends for (a) ZRE 1, (b) ZRE 2, and (c) ZRE 3, respectively. The distance is calculated between the ends of two monomers of the ZTA protein from the ZTA – DNA complex over the 200 ns unbiased MD simulation trajectories. The core motifs are shown in blue colour and the core motif with flanking ends in red colour.

### Structural stability analysis of dsDNA at the binding domain Minor groove analysis

Since the ZTA protein binds with dsDNA at the major and minor groove regions, any changes in the lengths of these grooves can potentially impact the binding and structural stability of the ZTA–dsDNA complex^38^. To understand the effect of adding flanking ends to the core motif and its effect on the overall stability and binding affinity of the ZTA–dsDNA complex, we calculated the DNA minor groove length for the core motif with flanking ends across the three ZRE systems. The average minor groove length was calculated over 200 ns of unbiased MD simulation trajectories. In Figure 17, we present a schematic of an instantaneous snapshot showing structural deformation of dsDNA (representative of ZRE 3 for the core motif with flanking ends) after 200 ns of unbiased MD simulation for the ZTA–dsDNA complex. In Figure 17 (a), we show the equilibrium structure of dsDNA highlighting a well-defined minor and major groove. The minor groove is marked by a blue dashed line, while the major groove is highlighted with a red dashed line. In Figure 17 (b), the dsDNA structure displays significant bending due to interaction with ZTA, resulting in an increase in minor groove length. Figure 18 illustrates the variation in minor groove length over time, and Table 5 provides quantitative estimates of the minor groove length for the three ZRE systems with flanking ends. The average minor groove length of the dsDNA system is approximately 12 Å. Upon adding the flanking sequences to the core motif, we observe average minor groove lengths of 11.56 (±1.29) Å for ZRE 1 (Figure 18 (a)), 12.29 (±1.13) Å for ZRE 2 (Figure 18 (b)), and 12.79 (±0.71) Å for ZRE 3 (Figure 18 (c)). This conformational change highlights the dynamic nature of dsDNA under the influence of ZTA protein. The increase in minor groove length and decrease in major groove length make the dsDNA more flexible, providing ZTA protein with greater conformational space to form the ZTA–dsDNA complex. Thus, the addition of DNA flanking ends induces structural changes in dsDNA that facilitate improved binding at the ZTA binding domain. Additionally, the FES values in Figure 7 and Table 4 show a reduction in free energy values, which supports this enhanced binding between ZTA and dsDNA.

**Figure 17:**
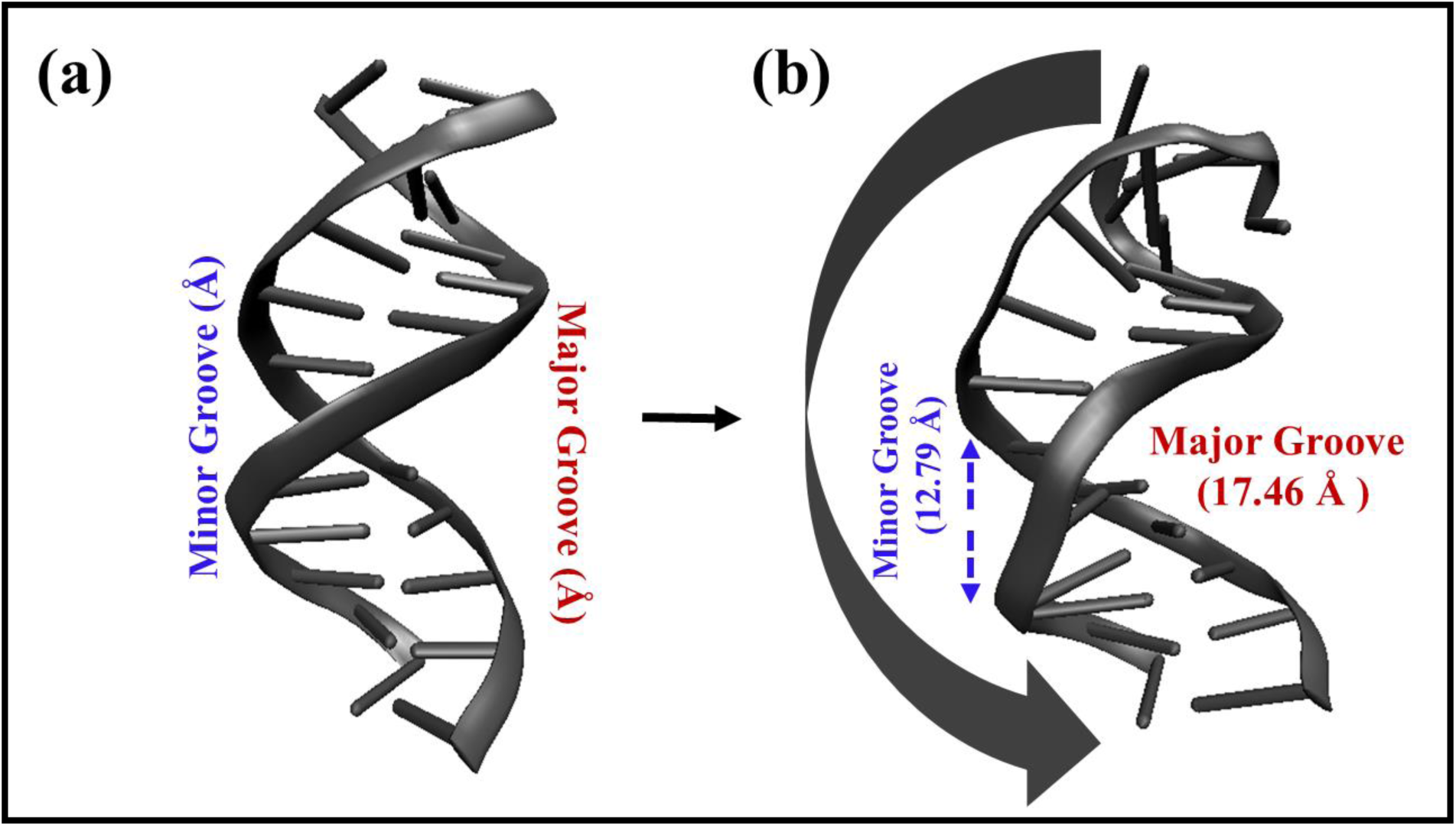
Instantaneous snapshot demonstrates the structural deformation of dsDNA (ZRE 3 for core motif with flanking ends) after 200 ns of unbiased MD simulation for the ZTA – DNA complex. In Figure (a) we show representative schematic of the minor and major grooves from the crystal structure of dsDNA. The minor groove is marked with blue dashed line, while the major groove is highlighted with a red dashed line. In figure (b) the dsDNA structure exhibits significant bending because of the interaction with ZTA with an increment in minor groove length. This conformational change underscores the dynamic nature of the dsDNA under the influence of ZTA protein.

**Figure 18:**
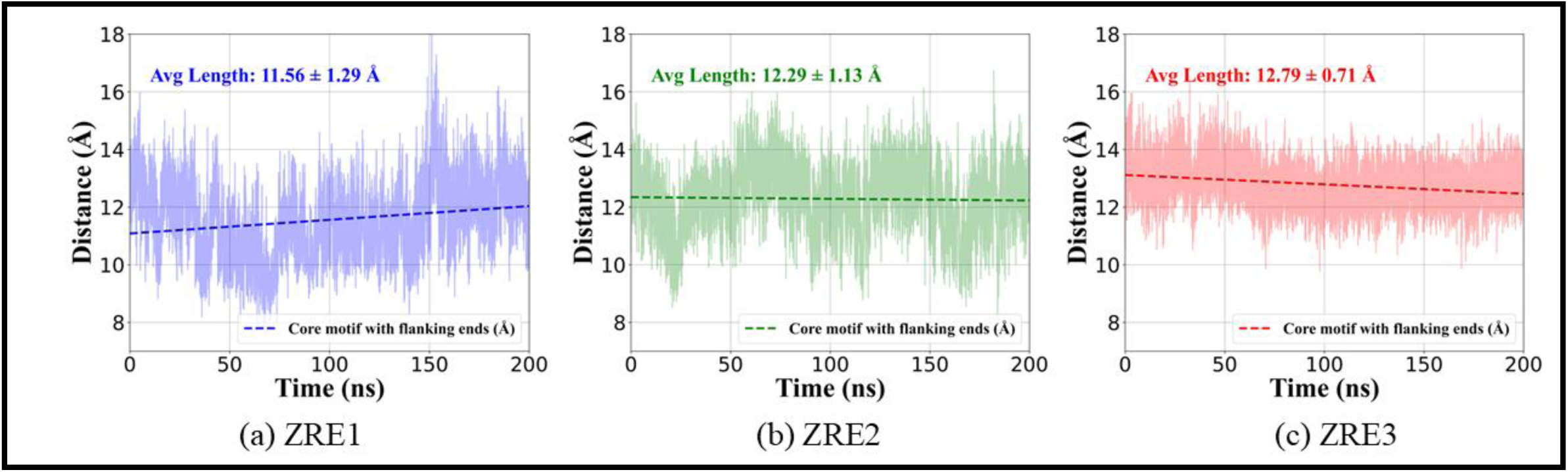
Distance (average minor groove length) vs time plot for the core motif with flanking ends for (a) ZRE 1, (b) ZRE 2, and (c) ZRE 3, respectively. The average minor groove length is calculated over the 200 ns unbiased MD simulation trajectories.

**Table 5:** Average minor groove lengths for three ZRE systems for the core motif with flanking ends sequences.

| DNA | ZRE sequences | Average minor groove length ± error bar (nm) |
| --- | --- | --- |
| core motif with flanking ends | ZRE1 | $\sim 12.29 \pm 1.13$ |
| | ZRE2 | $\sim 11.56 \pm 1.29$ |
| | ZRE3 | $\sim 12.79 \pm 0.71$ |

### H-bond analysis between DNA strands

Since the addition of DNA flanking ends affects the lengths of the major and minor grooves, it is expected to also alter interactions between the two DNA strands. As two single-stranded DNA strands primarily form hydrogen bonds to create dsDNA, we further calculated the number of hydrogen bonds between the two dsDNA strands. These hydrogen bonds were calculated from unbiased MD simulation trajectories. In Figure 19, we present the probability of hydrogen bonds between the two DNA strands (chain 1 and chain 2) in a ZTA protein–DNA complex. Figure 19 (a) shows the number of hydrogen bonds for the core motif systems, while Figure 19 (b) illustrates those for the core motif with flanking ends for ZRE 1 (blue), ZRE 2 (green), and ZRE 3 (red), respectively. As seen in Figure 19 (a), for ZRE 1 and ZRE 2 (core motif), the number of hydrogen bonds is approximately 8 with a probability of 0.12, while for ZRE 3 (core motif), it is around 11 with a probability of 0.10. Therefore, the number of interstrand hydrogen bonds in the dsDNA for the core motif cases follows the trend ZRE 1 (∼ 8) ≈ ZRE 2 (∼8) < ZRE 3 (∼11). For the core motif with flanking ends, shown in Figure 19 (b), ZRE 1, ZRE 2, and ZRE 3 exhibit approximately 16 hydrogen bonds, each with a probability of around 0.09. Thus, adding flanking ends to the DNA core motif increases the number of interstrand hydrogen bonds to around 16, compared to ∼ 8-10 for the core motif systems. This increase provides greater stability to the overall ZTA-dsDNA complex.

**Figure 19:**
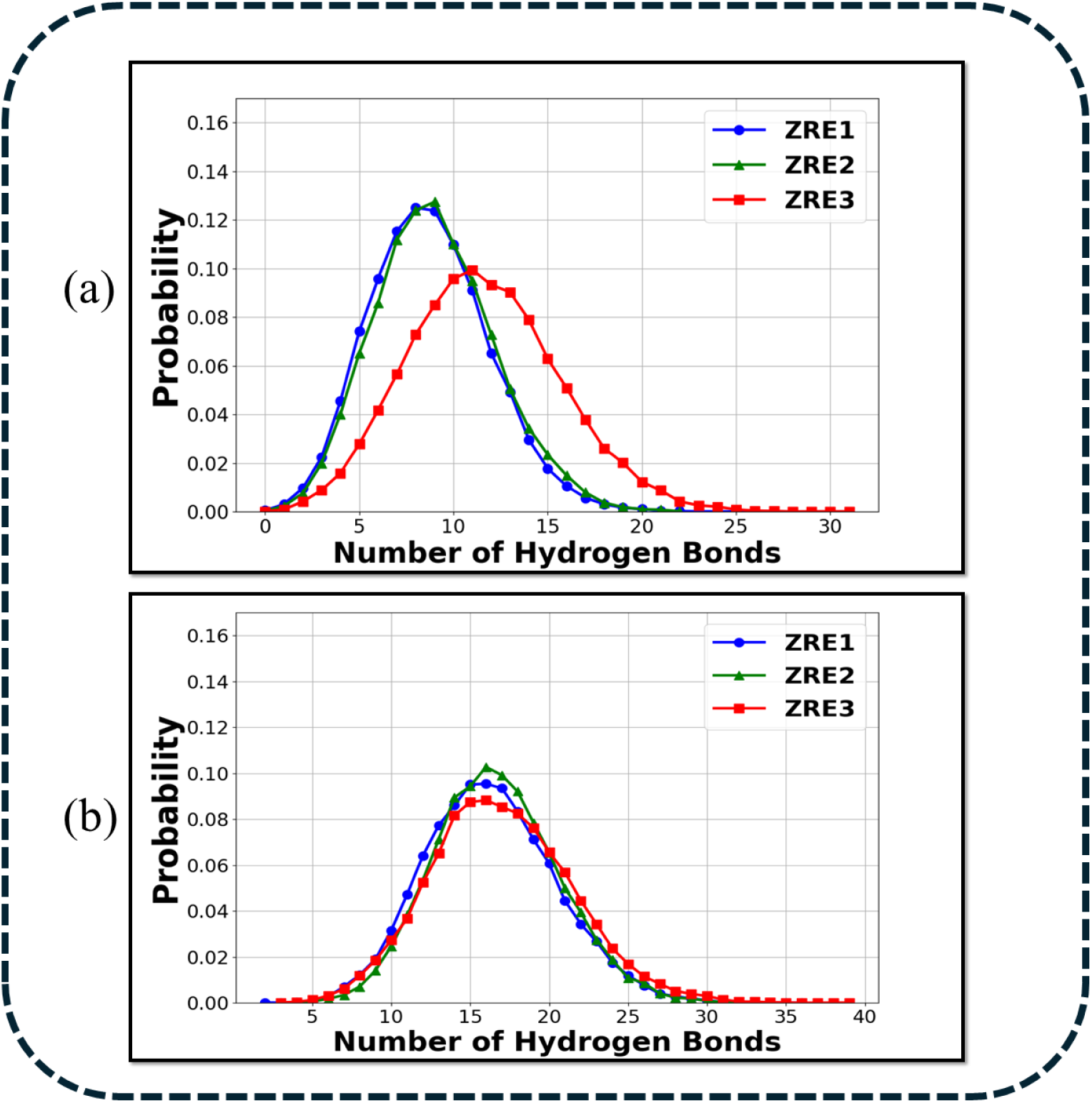
Probability of hydrogen bonds between the two DNA strands (chain 1 and chain 2) in a ZTA protein – DNA complex. Figure (a) core motif, and (b) core motif with flanking ends, for ZRE 1 (blue), ZRE 2 (green) and ZRE 3 (red) respectively. The hydrogen bonds are calculated from the unbiased MD simulation trajectories.

## Discussion

In this study, we employed all-atom molecular dynamics simulation combined with umbrella sampling technique to elucidate the structural stability and binding affinity of the ZTA transcription factor-dsDNA complex for the core motif and core motif with DNA flanking ends. We studied three ZRE systems (ZRE 1, ZRE 2, and ZRE 3), modelling each and performing 200 ns unbiased MD simulations. We calculated RMSD, RMSF and COM distances between ZTA and dsDNA from the unbiased MD simulation trajectories. These structural properties indicate that variations in DNA core motif sequences affect the binding affinity, and consequently the stability of ZTA with dsDNA. When we add flanking end sequences to the DNA core motif, the stability of the ZTA-dsDNA complex is enhanced for all three ZRE systems.

To understand the binding behaviour of this complex and to estimate quantitative binding affinity between ZTA and dsDNA, we calculated the free energy between ZTA and dsDNA using the COM–COM distance between ZTA and dsDNA groups as the reaction coordinate. Free energy values show that for the core motif cases, Δ*G* values range approximately from - 33 to -46 kcal/mol, whereas, with the addition of flanking ends to the core motif sequences, Δ*G* values lie in the range of -67 to -82 kcal/mol. Therefore, the addition of flanking ends to the core motif sequences significantly improves the stability of the dsDNA and ZTA TF complex, as seen in the substantially lower free energy values across all ZRE systems. Especially for ZRE 3, for the core motif with flanking ends, the lowest free energy value indicates optimal stability for the ZTA TF-dsDNA complex. To understand the basis of the structural stability and strong binding affinity between the ZTA TF and dsDNA complex, we calculated major interactions (anionic, van der Waals (VDW) and hydrophobic) that persisted for more than 90% of the simulation time. For the core motif case, the total number of interactions follows the trend: ZRE 2 > ZRE 1 > ZRE 3, and for the core motif with flanking ends, ZRE 3 > ZRE 2 > ZRE 1. We show that for the core motif with flanking ends, ZRE 3 has the highest number of interactions of various types, providing greater stability. Among the various interactions, van der Waals interactions emerge as the most dominant across all complexes. To identify specific protein residues that significantly contribute to these interactions and provide stability to the complex, we conducted a residue-level analysis. We observed that the amino acid arginine forms most interactions with dsDNA. Likewise, a nucleic acid-level analysis shows that, except for ZRE 1 (core motif), other ZRE systems, for both core motif and core motif with flanking ends, involve interactions formed by two, three, or all four bases (A, T, G, C) with protein residues. For ZRE 3 core motif with flanking ends, G nucleic bases play major role in forming interactions. Thus, free energy and total interaction counts show a good agreement for the ZRE 3, core motif with flanking ends, having lowest free energy and highest number of interactions. Additionally, the presence of a higher number of G interactions likely contribute to the stability, along with the overall interaction count. Hong et al.,^32^ previously showed that the presence of C and G nucleic bases in DNA enhances the stability of the ZTA–dsDNA complex. In our study, we observed a similar trend for both the core motif and core motif with flanking ends, with the CG base/pair numbers follow the order ZRE 3 > ZRE 2 > ZRE 1. This trend in C/G correlates well with the free energy values showing a good agreement. Beyond VDW and other interactions, we also found hydrogen bonds between the ZTA and dsDNA strands follows the order ZRE 1 (∼ 12) < ZRE 2 (∼18) ≤ ZRE 3 (∼18) for the core motif case, similar to the observed order in the free energy values (ΔG) for the core motif sequences: ZRE 1 ∼ -33.47 ± 0.41 kcal/mol, ZRE 2 ∼ -46.33 ± 0.36 kcal/mol, and ZRE 3 ∼ -46.04 ± 0.25 kcal/mol. For the core motif with flanking ends, the hydrogen bond count follows the trend ZRE 1 (∼ 19) ≤ ZRE 2 (∼18) < ZRE 3 (∼22) for the core motif with flanking ends. The trend in free energy values (ΔG) for the core motif with flanking ends is ZRE 1 ∼ -67.87 ± 0.40 kcal/mol, ZRE 2 ∼ -76.43 ± 0.50 kcal/mol, and ZRE 3∼ -82.06 ± 0.34 kcal/mol. Thus, a strong correlation is observed between hydrogen bonds and free energy, particularly for ZRE 3 with the core motif with flanking ends, where approximately 22 hydrogen bonds, align with the lowest ΔG value. This suggest that hydrogen bonding is a key factor contributing to the stability of the ZTA–dsDNA complex. For the dsDNA systems, in the core motif with flanking ends, minor groove lengths increase compared to the core motif systems. The increase in minor groove length coupled with a decrease in major groove length, makes dsDNA more flexible, providing ZTA greater conformational space to form the ZTA–dsDNA complex. Therefore, adding DNA flanking ends induces structural changes in dsDNA that facilitate improved binding at the ZTA binding domain. This conformational change highlights the dynamic nature of dsDNA under the influence of ZTA protein. Addition of flanking ends to the DNA core motif increases the number of interstrand hydrogen bonds to around 16, compared to ∼ 8-10 for the core motif systems. The higher number of hydrogen bonds supports the observed changes in minor and major groove lengths for dsDNA, enhancing the stability of the ZTA- dsDNA complex.

This finding further corroborates the increased stability and binding affinity observed in the free energy and structural stability results across all systems analysed. This comprehensive approach across multiple analyses clearly concludes that ZRE 3’s enhanced stability and binding affinity arise from specific interactions with C and G bases, both in core motif and core motif with flanking ends. This study highlights the substantial role of DNA sequence and structural context in modulating ZTA-DNA interaction dynamics within viral genomic regulation.

## Conclusion

This study investigated the structural stability and binding affinity of the ZTA transcription factor-dsDNA complex for the core motif and core motif with flanking ends using all-atom molecular dynamics simulation combined with umbrella sampling technique. When flanking sequences are added to the DNA core motif, the structural stability is enhanced for all three ZRE systems. Free energy calculations also show that addition of flanking ends sequences to the core motif lower the free energy values for all three ZRE systems. In terms of binding, we observe van der Waals (VDW) interaction to be the most dominant interaction across all the systems. The arginine residue plays the most important role in forming interactions with dsDNA, whereas for the nucleic acids, there was no specific preference except for ZRE 3 with the core motif with flanking ends, where G base plays the most dominant role. The CG base pair numbers also show a good correlation with free energy and stability for all the ZRE systems. Addition of flanking end sequences to the core motif affects the DNA groove lengths, increasing the minor groove length while decreasing the major groove length. This change facilitates conformational flexibility in DNA, providing ZTA with improved binding possibilities. The interstrand hydrogen bonds also increase for the core motif with flanking end sequences, further supporting the change in dsDNA groove lengths. Therefore, we observe that addition of flanking end sequences to the core motif enhances overall structural stability and lowers the free energy values for all three ZRE systems. Among the three ZRE systems, ZRE 3 with core motif and flanking end sequences, shows the lowest free energy value, the highest number of total interactions and hydrogen bonds, the highest CG base pair counts, an increased minor groove length, and the most DNA interstrand hydrogen bonds. These combined properties make the ZRE 3 core motif with flanking end sequences the most stable system. These findings underscore the importance of natural sequences and flanking end sequence context in facilitating efficient and stable protein-DNA interactions. Future research should focus on designing targeted therapies that disrupt these key interactions, leveraging the detailed molecular insights gained from this study. This study highlights the substantial role of DNA sequence and structural context in modulating ZTA-DNA interaction dynamics within viral genomic regulation for the Epstein-Barr virus (EBV).

## Supplementary information

In the supplementary information we have provided the following: In Figure S1(a-f), for all the six systems we provided individual free energy plots separately and for each system there are 10 independent free energy plots; Figure S2, interaction fingerprint for core motif ZRE 1; Figure S3, interaction fingerprint for core motif ZRE 2; Figure S4, interaction fingerprint for core motif ZRE 3; Figure S5, interaction fingerprint for core motif with flanking ends for ZRE 1; Figure S6, interaction fingerprint for core motif with flanking ends ZRE 2; Figure S7, interaction fingerprint for core motif with flanking ends ZRE 3; In pg.(9-10) we added the modelling report of the crystal structure used this work, methodology of RMSD & RMSF calculation and Interaction criteria of each interaction type from ProLIF tool.

## Acknowledgments

BD and DP thank SRM University – AP for providing HPCC supercomputing facilities. DP thanks Science and Engineering Research Board (SERB), Government of India for supporting with the State University Research Excellence (SURE) project No SUR/2022/004576. BD thanks Anuj Kumar for help in the analysis.

## References

(1) Young, L. S.; Dawson, C. W. Epstein-Barr Virus and Nasopharyngeal Carcinoma. Chin. J. Cancer 2014. 10.5732/cjc.014.10197.

(2) Bonnet, M.; Guinebretiere, J.-M.; Kremmer, E.; Grunewald, V.; Benhamou, E.; Contesso, G.; Joab, I. Detection of Epstein-Barr Virus in Invasive Breast Cancers. JNCI J. Natl. Cancer Inst. 1999, 91 (16), 1376–1381. 10.1093/jnci/91.16.1376.

(3) Tokunaga, M.; Land, C. E.; Uemura, Y.; Tokudome, T.; Tanaka, S. Epstein-Barr Virus in Gastric Carcinoma. 1993, 143 (5).

(4) Negro, F. The Paradox of Epstein–Barr Virus-Associated Hepatitis. J. Hepatol. 2006, 44 (5), 839–841. 10.1016/j.jhep.2006.03.002.

(5) Sarwari, N. M.; Khoury, J. D.; Hernandez, C. M. R. Chronic Epstein Barr Virus Infection Leading to Classical Hodgkin Lymphoma. BMC Hematol. 2016, 16 (1), 19. 10.1186/s12878-016-0059-3.

(6) Khan, G.; Hashim, M. J. Global Burden of Deaths from Epstein-Barr Virus Attributable Malignancies 1990-2010. Infect. Agent. Cancer 2014, 9 (1), 38. 10.1186/1750-9378-9-38.

(7). Ebell, M. H. Epstein-Barr Virus Infectious Mononucleosis. 2004, *70* (7).

(8) Lässer, C.; Seyed Alikhani, V.; Ekström, K.; Eldh, M.; Torregrosa Paredes, P.; Bossios, A.; Sjöstrand, M.; Gabrielsson, S.; Lötvall, J.; Valadi, H. Human Saliva, Plasma and Breast Milk Exosomes Contain RNA: Uptake by Macrophages. J. Transl. Med. 2011, 9 (1), 9. 10.1186/1479-5876-9-9.

(9) Luderer, R.; Kok, M.; Niesters, H. G. M.; Schuurman, R.; Weerdt, O.; Thijsen, S. F. T. Real-Time Epstein-Barr Virus PCR for the Diagnosis of Primary EBV Infections and EBV Reactivation. Mol. Diagn. 2005, 9 (4), 195–200. 10.1007/BF03260091.

(10) Braz-Silva, P. H.; De Rezende, N. P. M.; Ortega, K. L.; De Macedo Santos, R. T.; De Magalhães, M. H. C. G. Detection of the Epstein–Barr Virus (EBV) by In Situ Hybridization as Definitive Diagnosis of Hairy Leukoplakia. Head Neck Pathol. 2008, 2 (1), 19–24. 10.1007/s12105-007-0039-9.

(11) Hammerschmidt, W. The Epigenetic Life Cycle of Epstein–Barr Virus. In Epstein Barr Virus Volume 1; Münz, C., Ed.; Current Topics in Microbiology and Immunology; Springer International Publishing: Cham, 2015; Vol. 390, pp 103–117. 10.1007/978-3-319-22822-8_6.

(12) Kano, K.; Katayama, T.; Takeguchi, S.; Asanome, A.; Takahashi, K.; Saito, T.; Sawada, J.; Saito, M.; Anei, R.; Kamada, K.; Miyokawa, N.; Nishihara, H.; Hasebe, N. Biopsy-proven Case of Epstein–Barr Virus (EBV)-associated Vasculitis of the Central Nervous System. Neuropathology 2017, 37 (3), 259–264. 10.1111/neup.12356.

(13) Kwok, H.; Wu, C. W.; Palser, A. L.; Kellam, P.; Sham, P. C.; Kwong, D. L. W.; Chiang, A. K. S. Genomic Diversity of Epstein-Barr Virus Genomes Isolated from Primary Nasopharyngeal Carcinoma Biopsy Samples. J. Virol. 2014, 88 (18), 10662–10672. 10.1128/JVI.01665-14.

(14) Martel-Renoir, D.; Grunewald, V.; Touitou, R.; Schwaab, G.; Joab, I. Qualitative Analysis of the Expression of Epstein--Barr Virus Lytic Genes in Nasopharyngeal Carcinoma Biopsies. J. Gen. Virol. 1995, 76 (6), 1401–1408. 10.1099/0022-1317-76-6-1401.

(15) Ladell, K.; Dorner, M.; Zauner, L.; Berger, C.; Zucol, F.; Bernasconi, M.; Niggli, F. K.; Speck, R. F.; Nadal, D. Immune Activation Suppresses Initiation of Lytic Epstein-Barr Virus Infection. Cell. Microbiol. 2007, 9 (8), 2055–2069. 10.1111/j.1462-5822.2007.00937.x.

(16) Murata, T. Regulation of Epstein–Barr Virus Reactivation from Latency. Microbiol. Immunol. 2014, 58 (6), 307–317. 10.1111/1348-0421.12155.

(17) Ramasubramanyan, S.; Osborn, K.; Al-Mohammad, R.; Naranjo Perez-Fernandez, I. B.; Zuo, J.; Balan, N.; Godfrey, A.; Patel, H.; Peters, G.; Rowe, M.; Jenner, R. G.; Sinclair, A. J. Epstein–Barr Virus Transcription Factor Zta Acts through Distal Regulatory Elements to Directly Control Cellular Gene Expression. Nucleic Acids Res. 2015, 43 (7), 3563–3577. 10.1093/nar/gkv212.

(18) Miller, G.; El-Guindy, A.; Countryman, J.; Ye, J.; Gradoville, L. Lytic Cycle Switches of Oncogenic Human Gammaherpesviruses1. In Advances in Cancer Research; Elsevier, 2007; Vol. 97, pp 81–109. 10.1016/S0065-230X(06)97004-3.

(19) Zhou, Y.; Heesom, K.; Osborn, K.; AlMohammed, R.; Sweet, S. M.; Sinclair, A. J. Identifying the Cellular Interactome of Epstein-Barr Virus Lytic Regulator Zta Reveals Cellular Targets Contributing to Viral Replication. J. Virol. 2020, 94 (3), e00927–19. 10.1128/JVI.00927-19.

(20) Zhao, M.; Nanbo, A.; Becnel, D.; Qin, Z.; Morris, G. F.; Li, L.; Lin, Z. Ubiquitin Modification of the Epstein-Barr Virus Immediate Early Transactivator Zta. J. Virol. 2020, 94 (22), e01298–20. 10.1128/JVI.01298-20.

(21) Chen, C.; Li, D.; Guo, N. Regulation of Cellular and Viral Protein Expression by the Epstein-Barr Virus Transcriptional Regulator Zta: Implications for Therapy of EBV Associated Tumors. Cancer Biol. Ther. 2009, 8 (11), 987–995. 10.4161/cbt.8.11.8369.

(22) Flemington, E. K.; Borras, A. M.; Lytle, J. P.; Speck, S. H. Characterization of the Epstein-Barr Virus BZLF1 Protein Transactivation Domain. J. Virol. 1992, 66 (2), 922–929. 10.1128/jvi.66.2.922-929.1992.

(23) Petosa, C.; Morand, P.; Baudin, F.; Moulin, M.; Artero, J.-B.; Müller, C. W. Structural Basis of Lytic Cycle Activation by the Epstein-Barr Virus ZEBRA Protein. Mol. Cell 2006, 21 (4), 565–572. 10.1016/j.molcel.2006.01.006.

(24) Sayers, E. W.; Bolton, E. E.; Brister, J. R.; Canese, K.; Chan, J.; Comeau, D. C.; Connor, R.; Funk, K.; Kelly, C.; Kim, S.; Madej, T.; Marchler-Bauer, A.; Lanczycki, C.; Lathrop, S.; Lu, Z.; Thibaud-Nissen, F.; Murphy, T.; Phan, L.; Skripchenko, Y.; Tse, T.; Wang, J.; Williams, R.; Trawick, B. W.; Pruitt, K. D.; Sherry, S. T. Database Resources of the National Center for Biotechnology Information. Nucleic Acids Res. 2022, 50 (D1), D20–D26. 10.1093/nar/gkab1112.

(25) Bhende, P. M.; Seaman, W. T.; Delecluse, H.-J.; Kenney, S. C. The EBV Lytic Switch Protein, Z, Preferentially Binds to and Activates the Methylated Viral Genome. Nat. Genet. 2004, 36 (10), 1099–1104. 10.1038/ng1424.

(26) Almohammed, R.; Osborn, K.; Ramasubramanyan, S.; Perez-Fernandez, I. B. N.; Godfrey, A.; Mancini, E. J.; Sinclair, A. J. Mechanism of Activation of the BNLF2a Immune Evasion Gene of Epstein-Barr Virus by Zta. J. Gen. Virol. 2018, 99 (6), 805–817. 10.1099/jgv.0.001056.

(27) Etheve, L.; Martin, J.; Lavery, R. Dynamics and Recognition within a Protein–DNA Complex: A Molecular Dynamics Study of the SKN-1/DNA Interaction. Nucleic Acids Res. 2016, 44 (3), 1440–1448. 10.1093/nar/gkv1511.

(28) Tan, C.; Terakawa, T.; Takada, S. Dynamic Coupling among Protein Binding, Sliding, and DNA Bending Revealed by Molecular Dynamics. J. Am. Chem. Soc. 2016, 138 (27), 8512–8522. 10.1021/jacs.6b03729.

(29) Etheve, L.; Martin, J.; Lavery, R. Protein–DNA Interfaces: A Molecular Dynamics Analysis of Time-Dependent Recognition Processes for Three Transcription Factors. Nucleic Acids Res. 2016, gkw841. 10.1093/nar/gkw841.

(30) Bernaudat, F.; Gustems, M.; Günther, J.; Oliva, M. F.; Buschle, A.; Göbel, C.; Pagniez, P.; Lupo, J.; Signor, L.; Müller, C. W.; Morand, P.; Sattler, M.; Hammerschmidt, W.; Petosa, C. Structural Basis of DNA Methylation-Dependent Site Selectivity of the Epstein–Barr Virus Lytic Switch Protein ZEBRA/Zta/BZLF1. Nucleic Acids Res. 2022, 50 (1), 490–511. 10.1093/nar/gkab1183.

(31) Yu, K.-P.; Heston, L.; Park, R.; Ding, Z.; Wang’ondu, R.; Delecluse, H.-J.; Miller, G. Latency of Epstein–Barr Virus Is Disrupted by Gain-of-Function Mutant Cellular AP-1 Proteins That Preferentially Bind Methylated DNA. Proc. Natl. Acad. Sci. 2013, 110 (20), 8176–8181. 10.1073/pnas.1301577110.

(32) Hong, S.; Wang, D.; Horton, J. R.; Zhang, X.; Speck, S. H.; Blumenthal, R. M.; Cheng, X. Methyl-Dependent and Spatial-Specific DNA Recognition by the Orthologous Transcription Factors Human AP-1 and Epstein-Barr Virus Zta. Nucleic Acids Res. 2017, 45 (5), 2503–2515. 10.1093/nar/gkx057.

(33) El-Guindy, A.; Ghiassi-Nejad, M.; Golden, S.; Delecluse, H.-J.; Miller, G. Essential Role of Rta in Lytic DNA Replication of Epstein-Barr Virus. J. Virol. 2013, 87 (1), 208–223. 10.1128/JVI.01995-12.

(34) Heston, L.; El-Guindy, A.; Countryman, J.; Dela Cruz, C.; Delecluse, H.-J.; Miller, G. Amino Acids in the Basic Domain of Epstein-Barr Virus ZEBRA Protein Play Distinct Roles in DNA Binding, Activation of Early Lytic Gene Expression, and Promotion of Viral DNA Replication. J. Virol. 2006, 80 (18), 9115–9133. 10.1128/JVI.00909-06.

(35) Ray, S.; Tillo, D.; Assad, N.; Ufot, A.; Deppmann, C.; Durell, S. R.; Porollo, A.; Vinson, C. Replacing C189 in the bZIP Domain of Zta with S, T, V, or A Changes DNA Binding Specificity to Four Types of Double-Stranded DNA. Biochem. Biophys. Res. Commun. 2018, 501 (4), 905–912. 10.1016/j.bbrc.2018.05.080.

(36) Ray, S.; Tillo, D.; Assad, N.; Ufot, A.; Porollo, A.; Durell, S. R.; Vinson, C. Altering the Double-Stranded DNA Specificity of the bZIP Domain of Zta with Site-Directed Mutagenesis at N182. ACS Omega 2022, 7 (1), 129–139. 10.1021/acsomega.1c04148.

(37) Park, R.; Heston, L.; Shedd, D.; Delecluse, H.-J.; Miller, G. Mutations of Amino Acids in the DNA-Recognition Domain of Epstein–Barr Virus ZEBRA Protein Alter Its Sub-Nuclear Localization and Affect Formation of Replication Compartments. Virology 2008, 382 (2), 145–162. 10.1016/j.virol.2008.09.009.

(38) Yella, V. R.; Bhimsaria, D.; Ghoshdastidar, D.; Rodríguez-Martínez, J. A.; Ansari, A. Z.; Bansal, M. Flexibility and Structure of Flanking DNA Impact Transcription Factor Affinity for Its Core Motif. Nucleic Acids Res. 2018, 46 (22), 11883–11897. 10.1093/nar/gky1057.

(39) Vasumathi, V.; Pramanik, D.; Sood, A. K.; Maiti, P. K. Structure of a Carbon Nanotube– Dendrimer Composite. Soft Matter 2013, 9 (4), 1372–1380. 10.1039/C2SM26804C.

(40) Gelpi, J.; Hospital, A.; Goñi, R.; Orozco, M. Molecular Dynamics Simulations: Advances and Applications. Adv. Appl. Bioinforma. Chem. 2015, 37. 10.2147/AABC.S70333.

(41) Pramanik, D.; Kanchi, S.; Ayappa, K. G.; Maiti, P. K. Dendrimers: A Novel Nanomaterial. In Computational Materials, Chemistry, and Biochemistry: From Bold Initiatives to the Last Mile; Shankar, S., Muller, R., Dunning, T., Chen, G. H., Eds.; Springer Series in Materials Science; Springer International Publishing: Cham, 2021; Vol. 284, pp 411–449. 10.1007/978-3-030-18778-1_19.

(42) Zhang, H.; Huang, F.; Gilbert, B.; Banfield, J. F. Molecular Dynamics Simulations, Thermodynamic Analysis, and Experimental Study of Phase Stability of Zinc Sulfide Nanoparticles. J. Phys. Chem. B 2003, 107 (47), 13051–13060. 10.1021/jp036108t.

(43) Pramanik, D.; Maiti, P. K. DNA-Assisted Dispersion of Carbon Nanotubes and Comparison with Other Dispersing Agents. ACS Appl. Mater. Interfaces 2017, 9 (40), 35287–35296. 10.1021/acsami.7b06751.

(44) Deublein, S.; Eckl, B.; Stoll, J.; Lishchuk, S. V.; Guevara-Carrion, G.; Glass, C. W.; Merker, T.; Bernreuther, M.; Hasse, H.; Vrabec, J. Ms2: A Molecular Simulation Tool for Thermodynamic Properties. Comput. Phys. Commun. 2011, 182 (11), 2350–2367. 10.1016/j.cpc.2011.04.026.

(45) Dutta, P.; Pramanik, D.; Singh, J. K. Phase Behavior of Pure PSPC and PEGylated Multicomponent Lipid and Their Interaction with Paclitaxel: An All-Atom MD Study. Langmuir 2021, 37 (34), 10259–10271. 10.1021/acs.langmuir.1c01049.

(46) Guterres, H.; Im, W. Improving Protein-Ligand Docking Results with High-Throughput Molecular Dynamics Simulations. J. Chem. Inf. Model. 2020, 60 (4), 2189–2198. 10.1021/acs.jcim.0c00057.

(47) Kumar, A.; Kukal, S.; Marepalli, A.; Kumar, S.; Govindarajan, S.; Pramanik, D. Probing the Molecular Interactions of A22 with Prokaryotic Actin MreB and Eukaryotic Actin: A Computational and Experimental Study. J. Phys. Chem. B 2024, acs.jpcb.4c02963. 10.1021/acs.jpcb.4c02963.

(48) Cole, D. J.; Tirado-Rives, J.; Jorgensen, W. L. Molecular Dynamics and Monte Carlo Simulations for Protein–Ligand Binding and Inhibitor Design. Biochim. Biophys. Acta BBA - Gen. Subj. 2015, 1850 (5), 966–971. 10.1016/j.bbagen.2014.08.018.

(49) Namsani, S.; Pramanik, D.; Khan, M. A.; Roy, S.; Singh, J. K. Metadynamics-Based Enhanced Sampling Protocol for Virtual Screening: Case Study for 3CLpro Protein for SARS-CoV-2. J. Biomol. Struct. Dyn. 2022, 40 (15), 7002–7017. 10.1080/07391102.2021.1892530.

(50) Smith, Z.; Pramanik, D.; Tsai, S.-T.; Tiwary, P. Multi-Dimensional Spectral Gap Optimization of Order Parameters (SGOOP) through Conditional Probability Factorization. J. Chem. Phys. 2018, 149 (23), 234105. 10.1063/1.5064856.

(51) Mollica, L.; Decherchi, S.; Zia, S. R.; Gaspari, R.; Cavalli, A.; Rocchia, W. Kinetics of Protein-Ligand Unbinding via Smoothed Potential Molecular Dynamics Simulations. Sci. Rep. 2015, 5 (1), 11539. 10.1038/srep11539.

(52) Ribeiro, J. M. L.; Tsai, S.-T.; Pramanik, D.; Wang, Y.; Tiwary, P. Kinetics of Ligand–Protein Dissociation from All-Atom Simulations: Are We There Yet? Biochemistry 2019, 58 (3), 156–165. 10.1021/acs.biochem.8b00977.

(53) Pramanik, D.; Maiti, P. K. Dendrimer Assisted Dispersion of Carbon Nanotubes: A Molecular Dynamics Study. Soft Matter 2016, 12 (41), 8512–8520. 10.1039/C6SM02015A.

(54) Tu, X.; Manohar, S.; Jagota, A.; Zheng, M. DNA Sequence Motifs for Structure-Specific Recognition and Separation of Carbon Nanotubes. Nature 2009, 460 (7252), 250–253. 10.1038/nature08116.

(55) Johnson, R. R.; Johnson, A. T. C.; Klein, M. L. Probing the Structure of DNA−Carbon Nanotube Hybrids with Molecular Dynamics. Nano Lett. 2008, 8 (1), 69–75. 10.1021/nl071909j.

(56) Johnson, R. R.; Kohlmeyer, A.; Johnson, A. T. C.; Klein, M. L. Free Energy Landscape of a DNA−Carbon Nanotube Hybrid Using Replica Exchange Molecular Dynamics. Nano Lett. 2009, 9 (2), 537–541. 10.1021/nl802645d.

(57) Torrie, G. M.; Valleau, J. P. Nonphysical Sampling Distributions in Monte Carlo Free-Energy Estimation: Umbrella Sampling. J. Comput. Phys. 1977, 23 (2), 187–199. 10.1016/0021-9991(77)90121-8.

(58) Ferrenberg, A. M.; Swendsen, R. H. Optimized Monte Carlo Data Analysis. Phys. Rev. Lett. 1989, 63 (12), 1195–1198. 10.1103/PhysRevLett.63.1195.

(59) Kumar, S.; Rosenberg, J. M.; Bouzida, D.; Swendsen, R. H.; Kollman, P. A. THE Weighted Histogram Analysis Method for Free-energy Calculations on Biomolecules. I. The Method. J. Comput. Chem. 1992, 13 (8), 1011–1021. 10.1002/jcc.540130812.

(60) Bienert, S.; Waterhouse, A.; de Beer, T. A. P.; Tauriello, G.; Studer, G.; Bordoli, L.; Schwede, T. The SWISS-MODEL Repository—New Features and Functionality. Nucleic Acids Res. 2017, 45 (D1), D313–D319. 10.1093/nar/gkw1132.

(61) Waterhouse, A.; Bertoni, M.; Bienert, S.; Studer, G.; Tauriello, G.; Gumienny, R.; Heer, F. T.; de Beer, T. A. P.; Rempfer, C.; Bordoli, L.; Lepore, R.; Schwede, T. SWISS-MODEL: Homology Modelling of Protein Structures and Complexes. Nucleic Acids Res. 2018, 46 (W1), W296– W303. 10.1093/nar/gky427.

(62) Pettersen, E. F.; Goddard, T. D.; Huang, C. C.; Couch, G. S.; Greenblatt, D. M.; Meng, E. C.; Ferrin, T. E. UCSF Chimera—A Visualization System for Exploratory Research and Analysis. J. Comput. Chem. 2004, 25 (13), 1605–1612. 10.1002/jcc.20084.

(63) Kemmish, H.; Fasnacht, M.; Yan, L. Fully Automated Antibody Structure Prediction Using BIOVIA Tools: Validation Study. PLOS ONE 2017, 12 (5), e0177923. 10.1371/journal.pone.0177923.

(64) Abraham, M. J.; Murtola, T.; Schulz, R.; Páll, S.; Smith, J. C.; Hess, B.; Lindahl, E. GROMACS: High Performance Molecular Simulations through Multi-Level Parallelism from Laptops to Supercomputers. SoftwareX 2015, 1–2, 19–25. 10.1016/j.softx.2015.06.001.

(65) Duan, Y.; Wu, C.; Chowdhury, S.; Lee, M. C.; Xiong, G.; Zhang, W.; Yang, R.; Cieplak, P.; Luo, R.; Lee, T.; Caldwell, J.; Wang, J.; Kollman, P. A Point-charge Force Field for Molecular Mechanics Simulations of Proteins Based on Condensed-phase Quantum Mechanical Calculations. J. Comput. Chem. 2003, 24 (16), 1999–2012. 10.1002/jcc.10349.

(66) Jorgensen, W. L.; Chandrasekhar, J.; Madura, J. D.; Impey, R. W.; Klein, M. L. Comparison of Simple Potential Functions for Simulating Liquid Water. J. Chem. Phys. 1983, 79 (2), 926–935. 10.1063/1.445869.

(67) Meza, J. C. Steepest Descent. WIREs Comput. Stat. 2010, 2 (6), 719–722. 10.1002/wics.117.

(68) Martoňák, R.; Laio, A.; Parrinello, M. Predicting Crystal Structures: The Parrinello-Rahman Method Revisited. Phys. Rev. Lett. 2003, 90 (7), 075503. 10.1103/PhysRevLett.90.075503.

(69) Van Gunsteren, W. F.; Berendsen, H. J. C. A Leap-Frog Algorithm for Stochastic Dynamics. Mol. Simul. 1988, 1 (3), 173–185. 10.1080/08927028808080941.

(70) Grubmüller, H.; Heller, H.; Windemuth, A.; Schulten, K. Generalized Verlet Algorithm for Efficient Molecular Dynamics Simulations with Long-Range Interactions. Mol. Simul. 1991, 6 (1– 3), 121–142. 10.1080/08927029108022142.

(71) Hess, B.; Bekker, H.; Berendsen, H. J. C.; Fraaije, J. G. E. M. LINCS: A Linear Constraint Solver for Molecular Simulations. J. Comput. Chem. 1997, 18 (12), 1463–1472. 10.1002/(SICI)1096-987X(199709)18:12<1463::AID-JCC4>3.0.CO;2-H.

(72) Essmann, U.; Perera, L.; Berkowitz, M. L.; Darden, T.; Lee, H.; Pedersen, L. G. A Smooth Particle Mesh Ewald Method. J. Chem. Phys. 1995, 103 (19), 8577–8593. 10.1063/1.470117.

(73) Bouysset, C.; Fiorucci, S. ProLIF: A Library to Encode Molecular Interactions as Fingerprints. J. Cheminformatics 2021, 13 (1), 72. 10.1186/s13321-021-00548-6.

(74) Hub, J. S.; De Groot, B. L.; Van Der Spoel, D. G_wham—A Free Weighted Histogram Analysis Implementation Including Robust Error and Autocorrelation Estimates. J. Chem. Theory Comput. 2010, 6 (12), 3713–3720. 10.1021/ct100494z.

(75) Bouysset, C.; Fiorucci, S. ProLIF: A Library to Encode Molecular Interactions as Fingerprints. J. Cheminformatics 2021, 13 (1), 72. 10.1186/s13321-021-00548-6.

