## Supplementary Information for "Influence of DNA sequences on thermodynamic and structural stability of ZTA transcription factor - DNA complex: An all-atom molecular dynamics study"

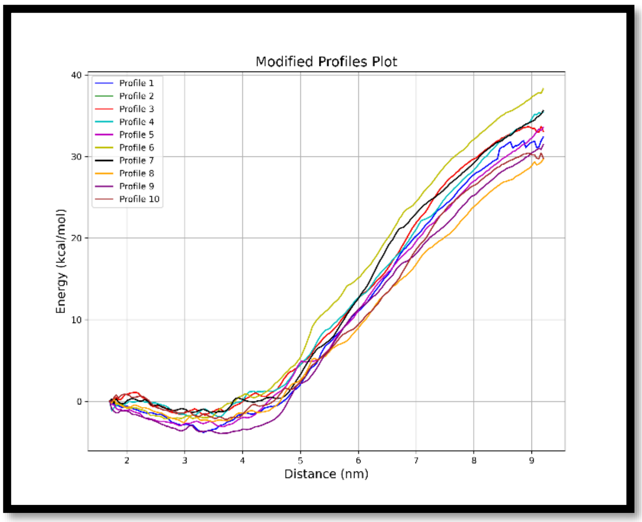
Modified Profiles Plot
Profile 1
Profile 2
Profile 3
Profile 4
Profile 5
Profile 6
Profile 7
Profile 8
Profile 9
Profile 10
Energy (kcal/mol)
40
30
20
10
0
2
3
4
5
6
7
8
9
Distance (nm)

(a)

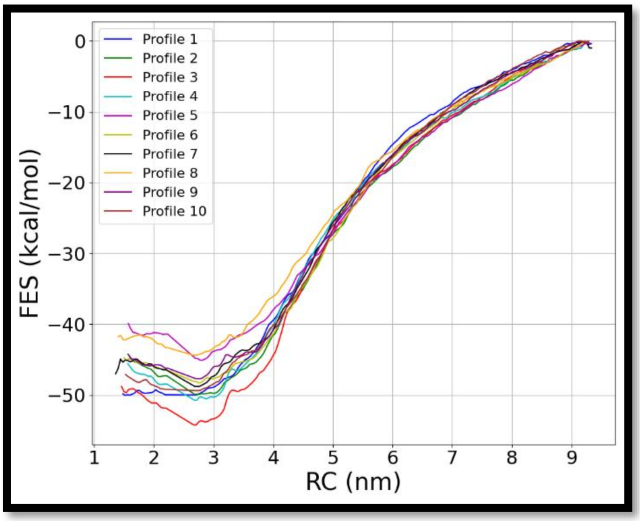
Profile 1
Profile 2
Profile 3
Profile 4
Profile 5
Profile 6
Profile 7
Profile 8
Profile 9
Profile 10
FES (kcal/mol)
0
−10
−20
−30
−40
−50
1
2
3
4
5
6
7
8
9
RC (nm)

(b)

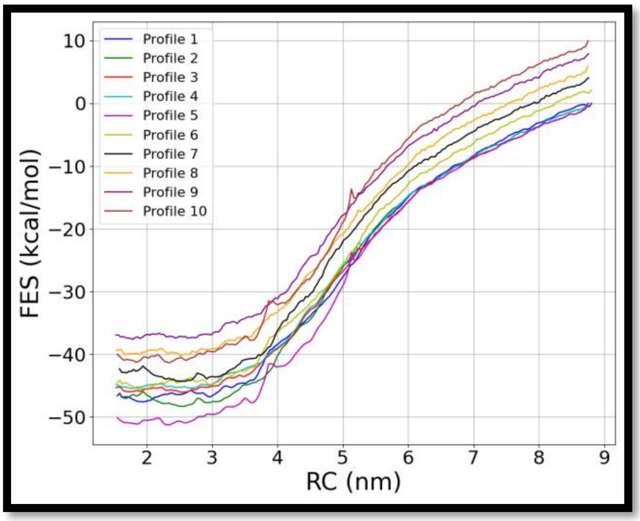
Profile 1
Profile 2
Profile 3
Profile 4
Profile 5
Profile 6
Profile 7
Profile 8
Profile 9
Profile 10
FES (kcal/mol)
10
0
−10
−20
−30
−40
−50
2
3
4
5
6
7
8
9
RC (nm)

(c)

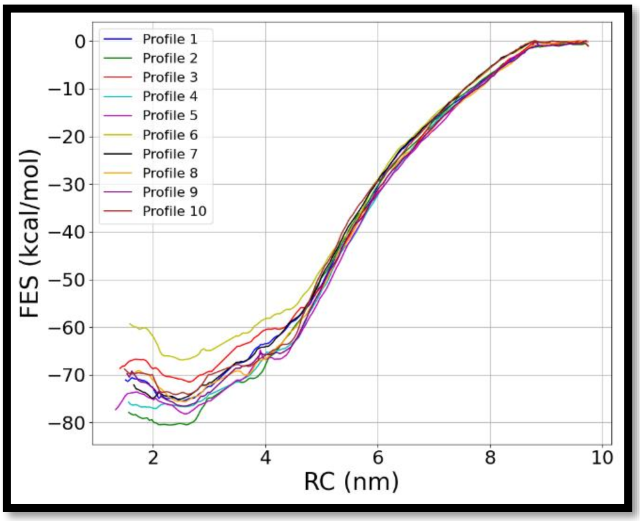
Profile 1
Profile 2
Profile 3
Profile 4
Profile 5
Profile 6
Profile 7
Profile 8
Profile 9
Profile 10
FES (kcal/mol)
0
−10
−20
−30
−40
−50
−60
−70
−80
2
4
6
8
10
RC (nm)

(d)

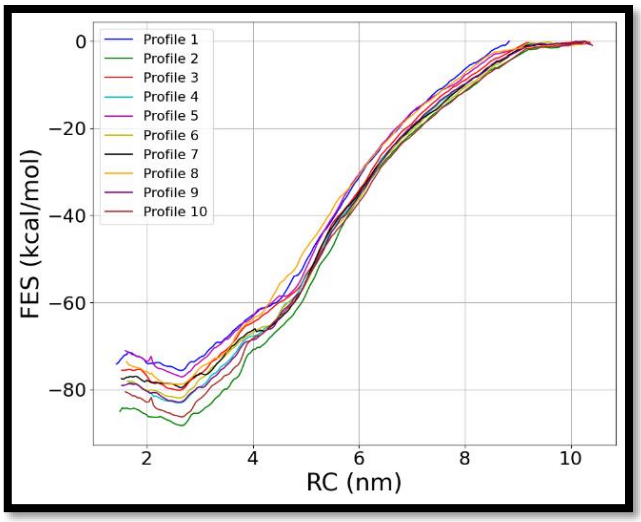
Profile 1
Profile 2
Profile 3
Profile 4
Profile 5
Profile 6
Profile 7
Profile 8
Profile 9
Profile 10
FES (kcal/mol)
0
−20
−40
−60
−80
2
4
6
8
10
RC (nm)

(e)

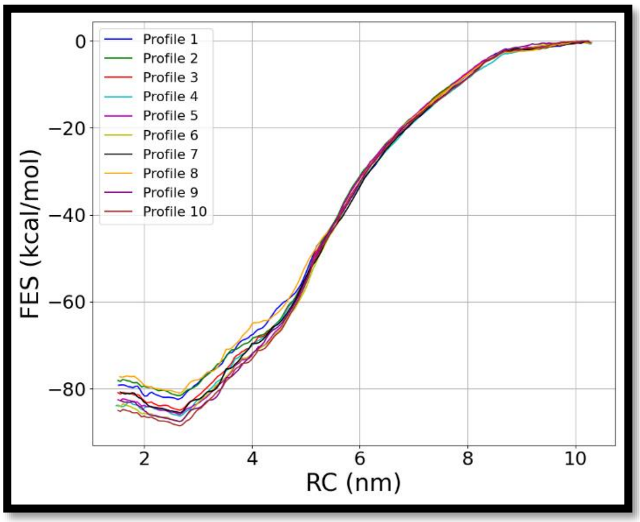
Profile 1
Profile 2
Profile 3
Profile 4
Profile 5
Profile 6
Profile 7
Profile 8
Profile 9
Profile 10
FES (kcal/mol)
0
−20
−40
−60
−80
2
4
6
8
10
RC (nm)

(f)

Figure S1: Free energy plots of 10 independent runs for each system, respectively (a) Zta TF with ZRE 1 (core motif), (b) Zta TF with ZRE 2 (core motif), (c) Zta TF with ZRE 3 (core motif), (d) Zta TF with ZRE 1 (core motif with flanking ends), (e) Zta TF with ZRE 2 (core motif with flanking ends), and (f) Zta TF with ZRE 3 (core motif with flanking ends).

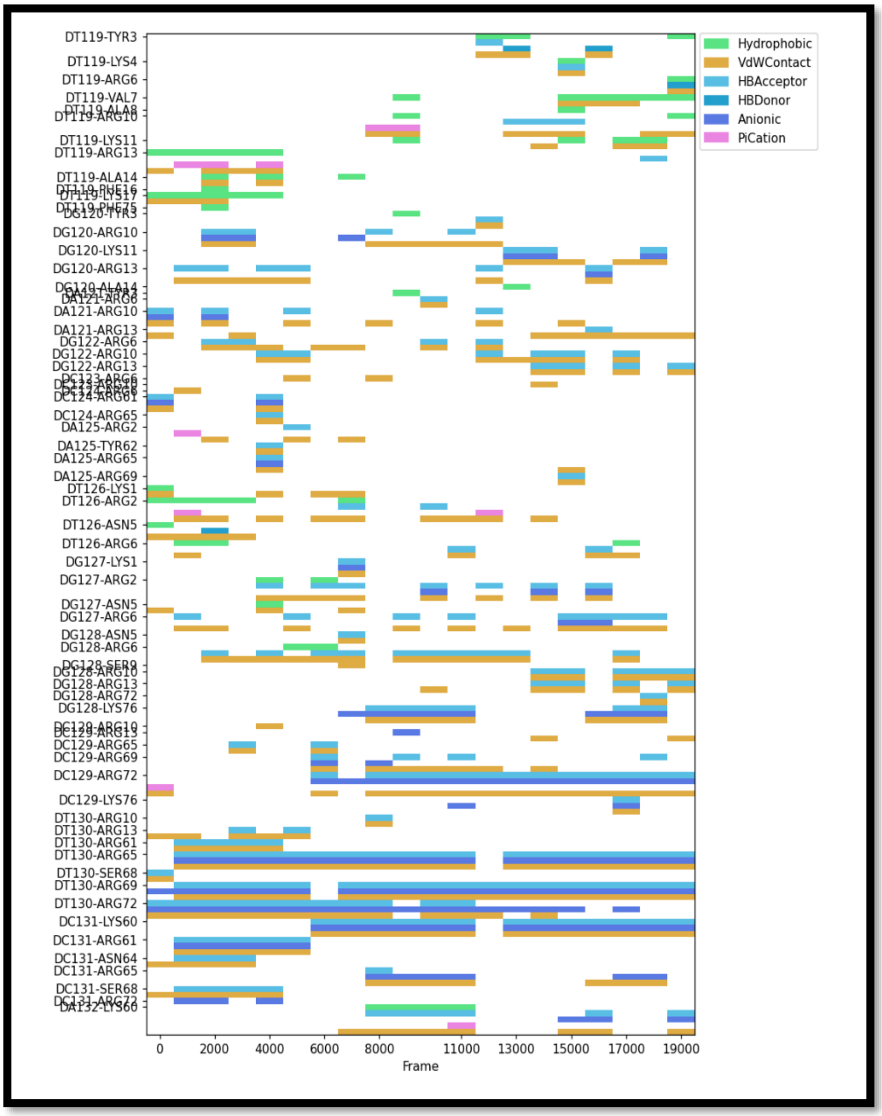


Figure S2: Interaction fingerprint for Zta TF with ZRE 1 dsDNA for the core motif case.

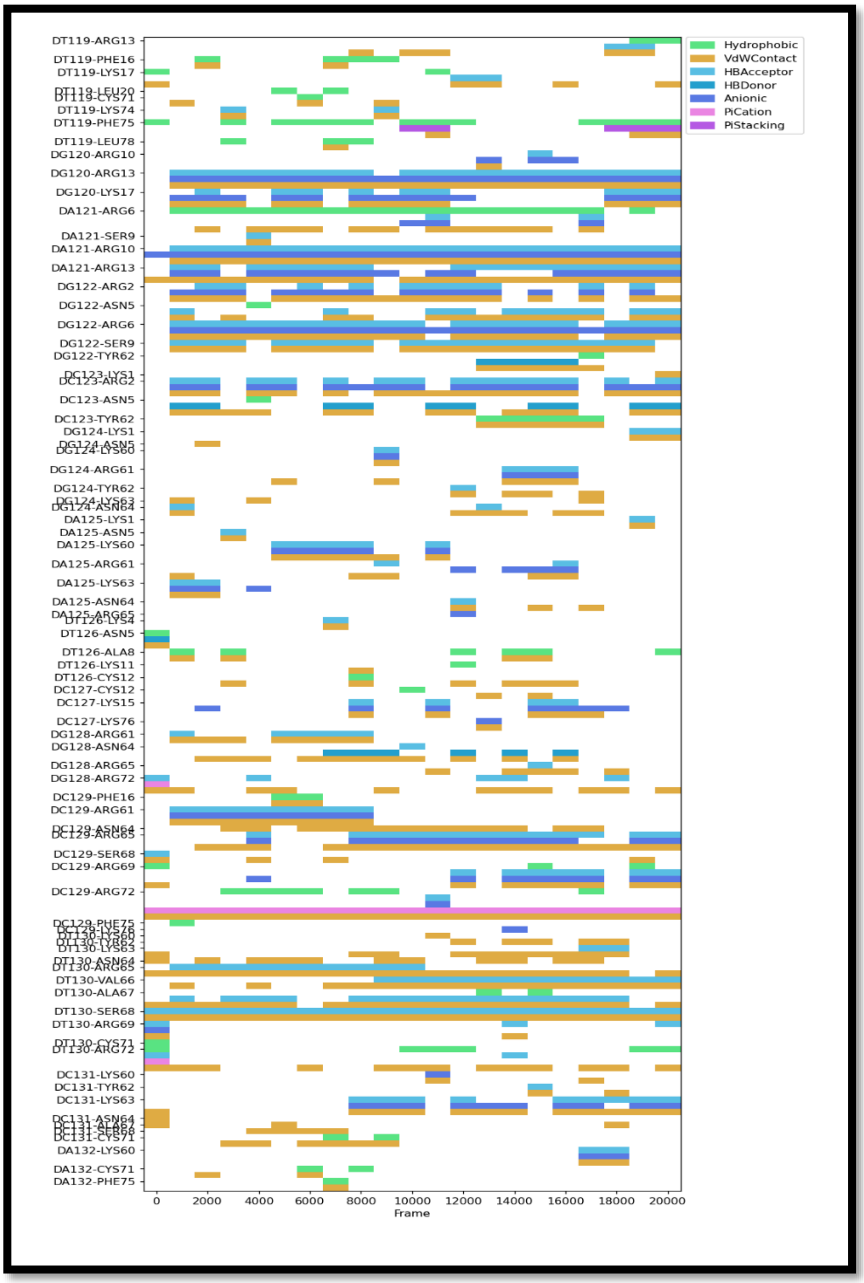


Figure S3: Interaction fingerprint for Zta TF with ZRE 2 dsDNA for the core motif case.

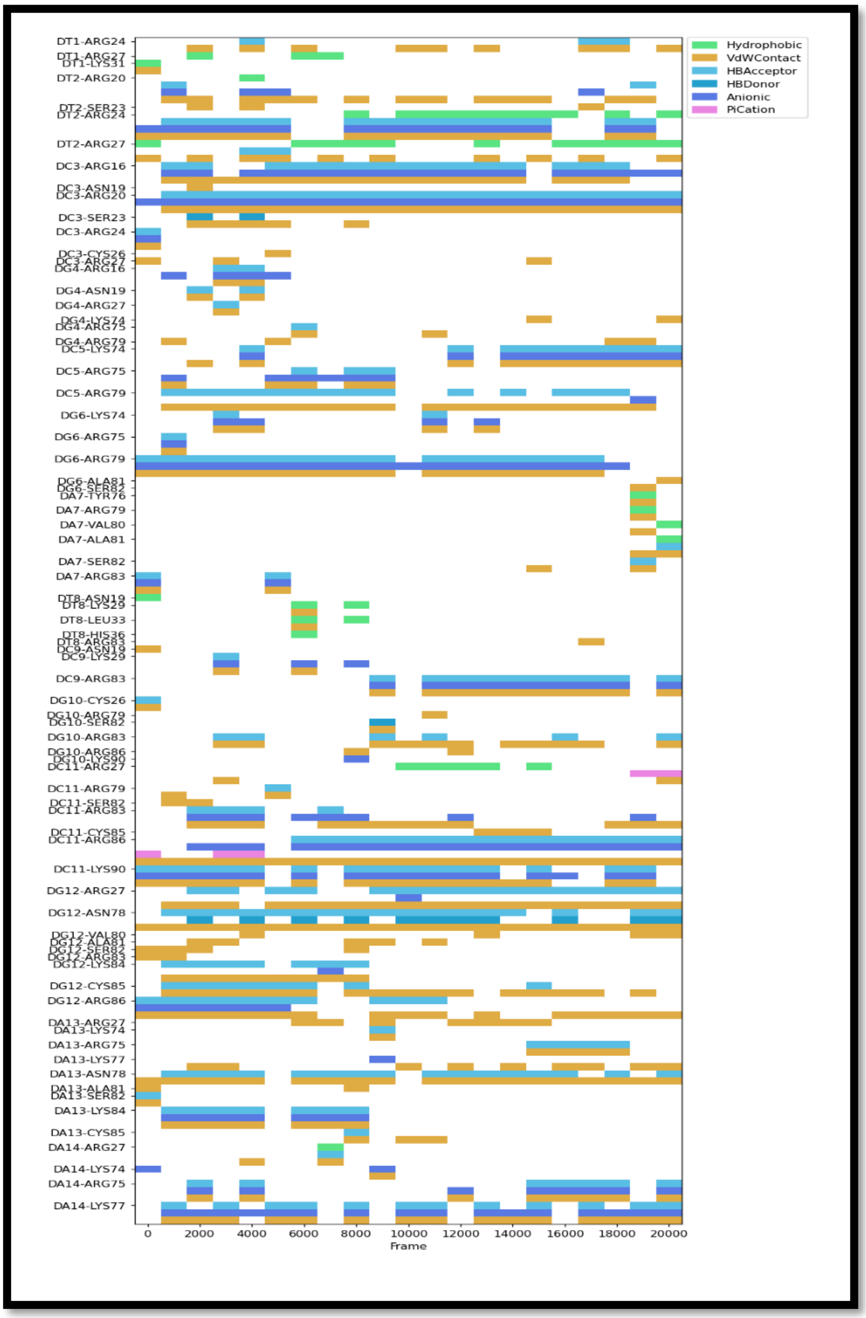


Figure S4: Interaction fingerprint for Zta TF with ZRE 3 dsDNA for the core motif case.

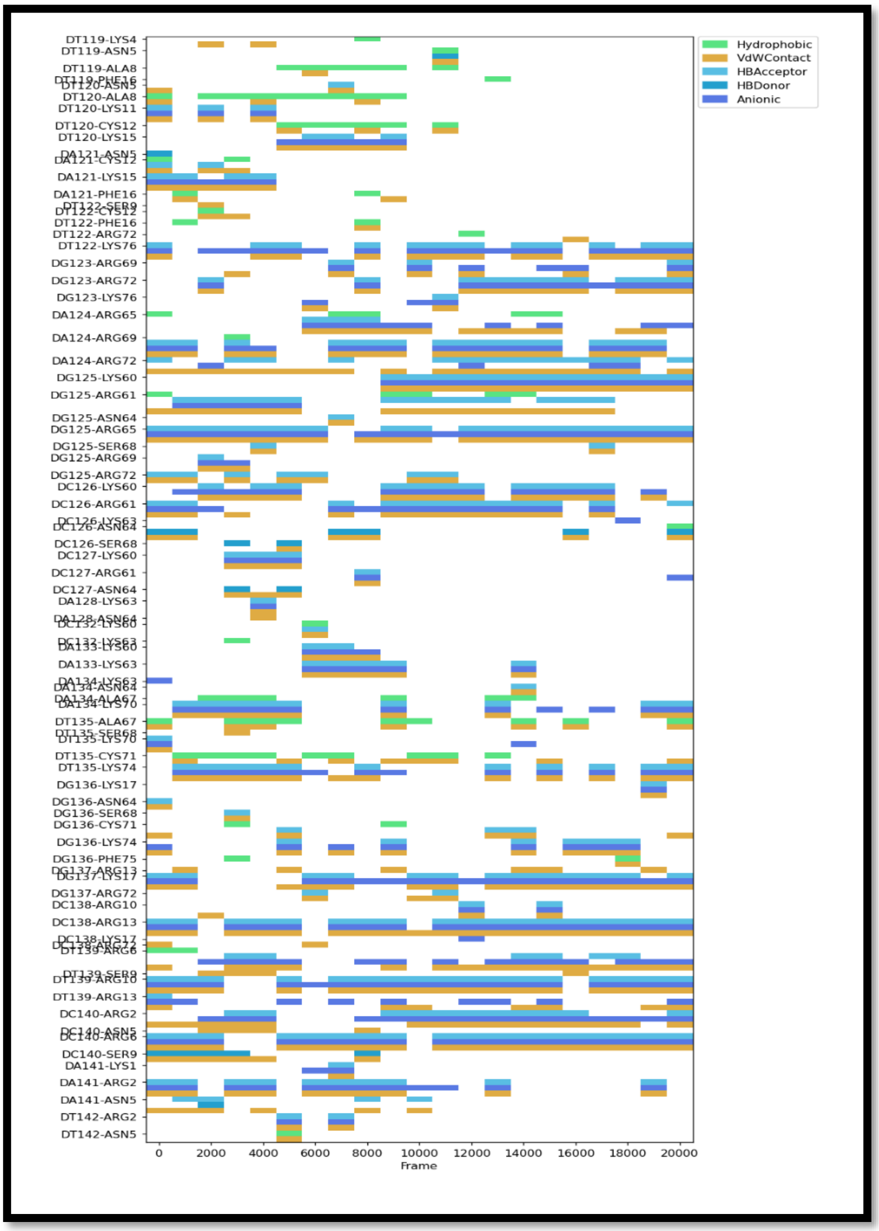


Figure S5: Interaction fingerprint for Zta TF with ZRE 1 dsDNA for the core motif with flanking ends.

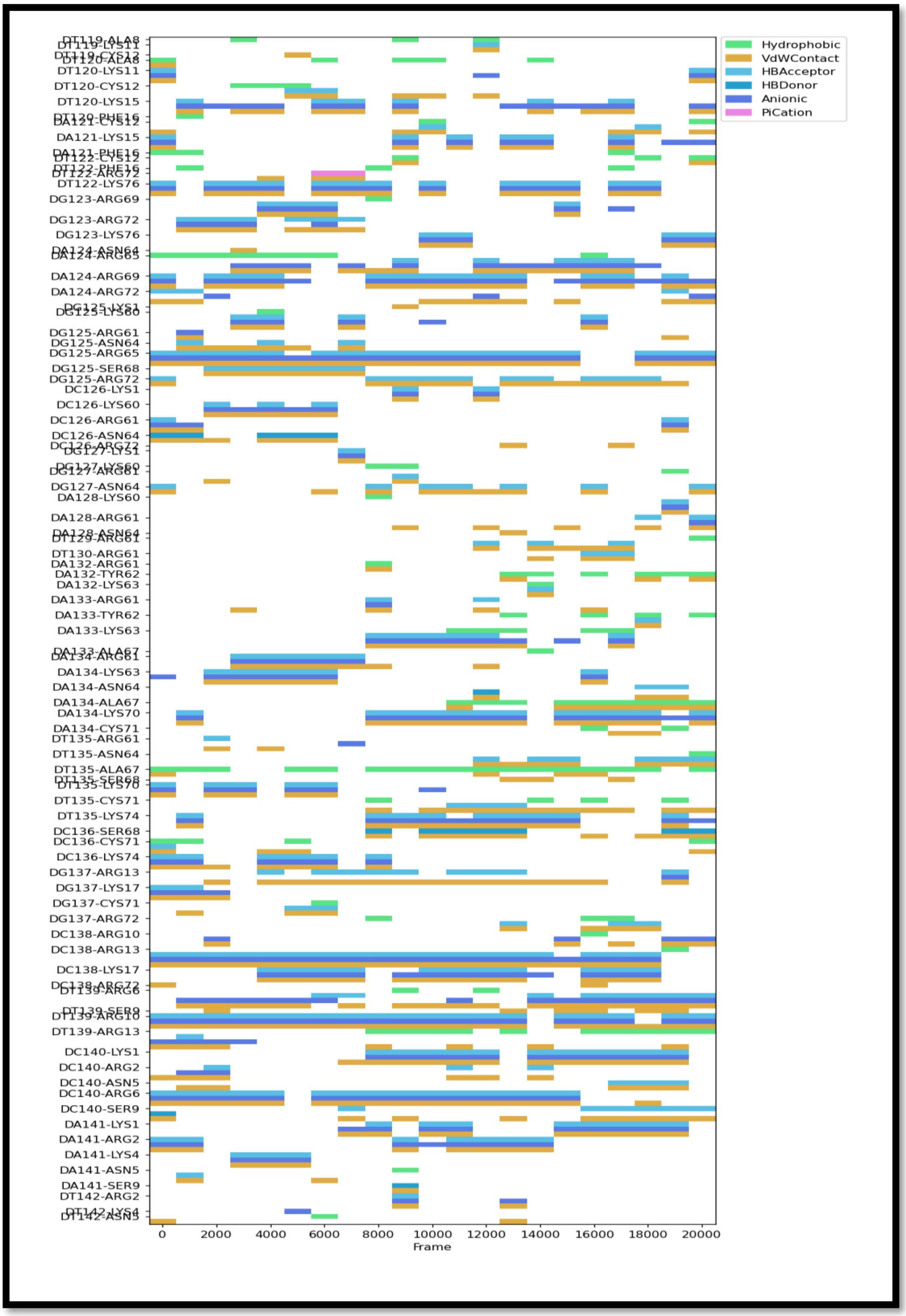


Figure S6: Interaction fingerprint for Zta TF with ZRE 2 dsDNA for the core motif with flanking ends.

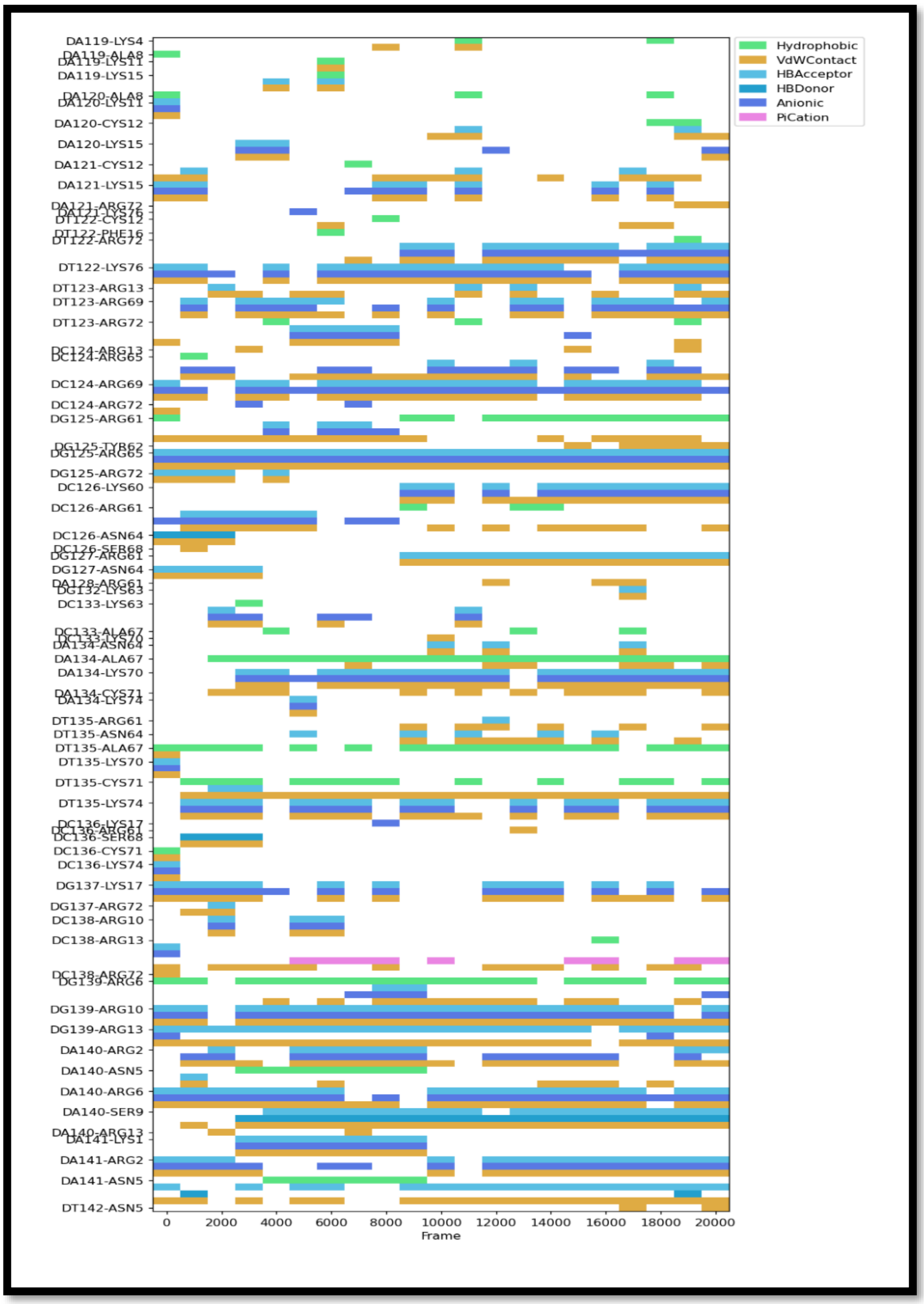


Figure S7: Interaction fingerprint for Zta TF with ZRE 3 dsDNA for the core motif with flanking ends.

**Zta-protein modeling:**

**Protein structure was downloaded using PDB ID:2C9N, and modeled using SWISS-Modeller,**

**Missing residue before modelling,**

```
REMARK 470 MISSING ATOM
REMARK 470 THE FOLLOWING RESIDUES HAVE MISSING ATOMS (M=MODEL NUMBER;
REMARK 470 RES=RESIDUE NAME; C=CHAIN IDENTIFIER; SSEQ=SEQUENCE NUMBER;
REMARK 470 I=INSERTION CODE):
REMARK 470   M RES CSSEQI  ATOMS
REMARK 470     MET A 174    CG   SD   CE
REMARK 470     LEU A 175    CG   CD1  CD2
REMARK 470     GLU A 176    CG   CD   OE1  OE2
REMARK 470     SER A 224    CB   OG
REMARK 470     GLU B 176    CB   CG   CD   OE1  OE2
REMARK 470     ILE B 177    CG2  CD1
REMARK 470     SER B 224    OG
REMARK 470     LEU B 225    CD1  CD2
REMARK 470      DC C  -8    P    OP1  OP2  O5'  C5'
REMARK 470      DG D  -7    P    OP1  OP2
REMARK 470     GLU E 176    CG   CD   OE1  OE2
REMARK 470     ARG E 179    CG   CD   NE   CZ   NH1  NH2
REMARK 470     VAL E 184    CG1  CG2
REMARK 470     LYS E 219    CD   CE   NZ
REMARK 470     ASP E 226    CG   OD1  OD2
REMARK 470     ASP E 228    CB   CG   OD1  OD2
REMARK 470     SER E 229    OG
REMARK 470     ARG E 233    CG   CD   NE   CZ   NH1  NH2
REMARK 470     THR E 234    CB   OG1  CG2
REMARK 470     ASP E 236    CG   OD1  OD2
REMARK 470     GLU F 176    CG   CD   OE1  OE2
REMARK 470     LYS F 181    CG   CD   CE   NZ
REMARK 470     SER F 186    OG
REMARK 470     ARG F 190    CG   CD   NE   CZ   NH1  NH2
REMARK 470     LEU F 197    CG   CD1  CD2
REMARK 470     TYR F 200    CG   CD1  CD2  CE1  CE2  CZ   OH
REMARK 470      DT G  -7    P    OP1  OP2  N1   C2   O2   N3
REMARK 470      DT G  -7    C4   O4   C5   C7   C6
```

### After modelling report,

```
REMARK   3 MODEL INFORMATION
REMARK   3  SMVERSN 2024-10.2
REMARK   3  ENGIN   PROMOD3
REMARK   3  VERSN   3.4.1
REMARK   3  OSTAT   homo-dimer
REMARK   3  OSRSN   PREDICTION
REMARK   3  QSPRD   0.470
REMARK   3  GMQE    0.10
REMARK   3  QMNV    4.3.1
REMARK   3  QMNDG   0.52
REMARK   3  MODT    FALSE
REMARK   3
REMARK   3 TEMPLATE 1
REMARK   3  PDBID   2c9n
REMARK   3  CHAIN   Z
REMARK   3  MMCIF   D
REMARK   3  PDBV    2024-10-04
REMARK   3  SMTLE   2c9n.1.D
REMARK   3  SMTLV   2024-10-09
REMARK   3  MTHD    X-RAY DIFFRACTION 3.30 A
REMARK   3  FOUND   HHblits
REMARK   3  GMQE    0.10
REMARK   3  SIM     0.60
REMARK   3  SID     100.00
REMARK   3  OSTAT   homo-dimer
REMARK   3  ALN D TRG MMDPNSTSEDVKFTPDPYQVPFVQAFDQATRVYQDLGGPSQAPLPCVLWPVLPEPLPQ
REMARK   3  ALN D TRG GQLTAYHVSTAPTGSWFSAPQPAPENAYQAYAAPQLFPVSDITQNQQTNQAGGEAPQP
REMARK   3  ALN D TRG GDNSTVQTAAAVVFACPGANQGQQLADIGVPQPAPVAAPARRTRKPQQPESLEECDSE
REMARK   3  ALN D TRG LEIKRYKNRVASRKCRAKFKQLLQHYREVAAAKSSENDRLRLLLKQMCPSLDVDSIIP
```

**RMSD & RMSF**

RMSD (root mean square deviation) of structure with respect to the initial reference structure can be calculated by least-square fitting the structure to the reference structure,

$$RMSD(t_1, t_2) = \left[\frac{1}{M}\sum_{i=1}^{N} m_i ||r_i(t_1) - r_i(t_2)||^2\right]^{\frac{1}{2}}$$

Where, $M = \sum_{i=1}^{N} m_i$ and $r_i(t)$ is the position of atom $i$ at time $t$ .

RMSF (root means quare fluctuation) is computed as the square root of the variance of the fluctuation around the average position,

$$\text{RMSF} = \left[\frac{1}{T}\sum_{t=1}^{T} (r_i(t) - \langle r_i \rangle)^2\right]^{\frac{1}{2}}$$

T is the total number of time points, $r_i(t)$ is the position of atom $i$ at time $t$ and $\langle r_i \rangle$ is the average position.

Hydrogen bond analysis (h-bond) between all possible donors D and acceptors A. To determine the exitance of and h-bond a geometrical criterion is used in gromacs[1].

Hydrogen bond (h-bond):

(h-bond) between all possible donors D and acceptors A. To determine the exitance of and h-bond a geometrical criterion is used in gromacs[1]

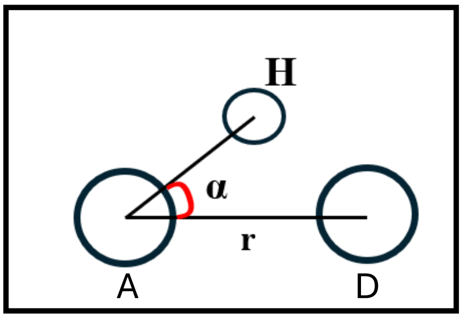


$r \leq r_{HB} = 0.35$ nm

$\alpha \leq \alpha_{HB} = 30°$

**Prolif interaction criteria:**

**Anionic interaction:**
Involves a negatively charged atom (anion) interacting with a [positively charged atom. The distance is less than or equal to 4.5 angs.

**Cationic:**
Involves a positively charged atom (cation) interacting with a negatively charged atom. The distance is less than or eual to 4.5 angs.

**Hydrophobic:**
Involves no-polar atoms interacts with distance of less thasn pr equal to 4.5 angs

**H-acceptor:**
A hydrogen bond forms between a hydrogen bond donor and an acceptor atom within less than or equal to 3.5 Å.

**H-donor:**
A dondor interacts with an acceptor atom, maintaining the same distance of less than or equal to 3.5 angs.
